# Quantitative analysis of genetic interactions in human cells from genome-wide CRISPR-Cas9 screens

**DOI:** 10.1101/2025.06.30.662330

**Authors:** Maximilian Billmann, Michael Costanzo, Mahfuzur Rahman, Katherine Chan, Amy Hin Yan Tong, Henry N. Ward, Arshia Z. Hassan, Xiang Zhang, Kevin R. Brown, Thomas Rohde, Angela H. Shaw, Catherine Ross, Jolanda van Leeuwen, Michael Aregger, Keith Lawson, Barbara Mair, Patricia Mero, Matej Usaj, Brenda Andrews, Charles Boone, Jason Moffat, Chad L. Myers

## Abstract

Genetic interaction (GI) networks in model organisms have revealed how combinations of genome variants can impact phenotypes. To advance efforts toward a reference human GI network, we developed the <u>q</u>uantitative <u>G</u>enetic Interaction (qGI) score, a method for precise GI measurement from genome-wide CRISPR-Cas9 screens in different query mutants constructed in a single human cell line. We found surprising prevalent systematic variation unrelated to GIs in CRISPR screen data, including both genomically linked effects and functionally coherent covariation. Leveraging ∼40 control screens in wild-type cells and half a billion differential fitness effect measurements, we developed a pipeline for CRISPR screen data processing and normalization to correct these artifacts and measure accurate, quantitative GIs. We also comprehensively characterized GI reproducibility by characterizing 4 – 5 biological replicates for ∼125,000 unique gene pairs. The qGI framework enables systematic identification of human GIs and provides broadly applicable strategies for analyzing context-specific CRISPR screen data.

## Main

Genetic interaction (GI) analysis is a powerful approach for systematic interrogation of gene function. A GI occurs when the combined perturbation of two genes produces an unexpected phenotype not predictable from the individual effects. Such interactions reflect functional relationships between genes^1^. For example, a negative GI occurs when double mutants exhibit a fitness defect greater than expected from the combined effects of the corresponding single mutants. Synthetic lethality represents an extreme negative GI, where each single mutant is viable and the double mutant is lethal. In some instances, synthetic lethal interactions can be therapeutically exploited, as demonstrated by the use of PARP1 inhibitors in BRCA1-mutant cancers^2,3^. Positive GIs reflect double mutants that grow better than expected from the combined single mutant phenotypes. Extreme positive interactions can reflect instances of genetic suppression, which can identify molecular mechanisms underlying protective variants in humans^1,4^. When GIs are systematically measured, the profile of a gene’s GIs captures its functional relationships and can be used to cluster genes in an unbiased fashion on a genome-scale^1^. In the yeast model system, exhaustive double mutant perturbation screens have been applied to successfully map a global network of functional relations between genes^5–7^. Similar yet less comprehensive endeavors in higher eukaryotes like *C. elegans* and *Drosophila* have used combinatorial RNAi^8–10^. Such systematic GI studies have relied heavily on the synergistic development of experimental and computational methods^7,11–18^.

In human cells, the systematic mapping of GIs has been facilitated by the development and application of CRISPR-based functional screening technologies^19–21^. Two primary strategies have emerged: combinatorial perturbation, where multiple gRNAs are co-expressed from a single vector^22–31^ and single-perturbation screens performed in cells carrying a stable pre-existing “query” (also called “anchor”) mutation^32–35^. Combinatorial systems differ mainly in how gRNA expression is controlled and typically induce acute perturbation of the targeted genes. These studies have thus far been limited in scale, targeting pairwise combinations of a few hundred genes within defined biological contexts, with the largest study to-date covering ∼150,000 gene pairs^36,37^. In contrast, CRISPR screens using genome-wide gRNA libraries represent an alternative approach for mapping GIs in cell lines that carry a stable mutation in a query gene of interest on a genome-wide scale^32–34,38^. Genes perturbed via gRNA libraries in the presence of a query mutation are referred to as library genes. As an alternative to the strategies inserting both mutations experimentally, several efforts have exploited genome-wide CRISPR screens in panels of cell lines that mimic a query mutation through naturally occurring mutations, DNA copy number or mRNA expression changes^39–43^, an approach that limits the search space for GIs to the genetic makeup of cell lines.

The effectiveness of these platforms, including the ability to scale up experiments, depends on the development of tailored computational pipelines. Data from combinatorial gRNA expression systems pose unique challenges, such as positional gRNA expression effects, and often require customized pre-processing workflows. Initial comparisons of computational strategies for analyzing dual perturbation data have begun to emerge^23,26,44–49^. In contrast, genome-scale query screens are often analyzed with methods adapted from standard CRISPR screens, which quantify how the abundance of cells carrying a certain perturbation changes over time^33,50–57^. These methods have proven highly effective for interpreting large-scale CRISPR screens across diverse cancer cell lines^53,54,58–60^. However, quantifying GIs requires measuring how CRISPR screen phenotypes change in the presence of a query mutation. Despite the availability of extensive genome-wide CRISPR screen datasets and refined analysis tools, comparable methods for systematic GI analysis remain limited, likely due to the scarcity of large datasets involving multiple genome-wide query screens.

Here, we introduce the <u>q</u>uantitative <u>G</u>enetic <u>I</u>nteraction (qGI) score, a method for precise identification of GIs from genome-wide CRISPR-Cas9 screens in isogenic human cell lines. Developed using hundreds of screens with the TKOv3 gRNA library^61^ in HAP1 cells, each harboring a defined loss-of-function mutation, the qGI score robustly measures interaction effect sizes and significance between library genes and the query gene. The qGI scoring pipeline fits fitness effects measured in a given HAP1 clone against a panel of HAP1 reference screens in a batch-aware fashion. We identify and correct unexpected chromosomal effects between isogenic HAP1 clones that otherwise introduce linkage-dependent false positive (FP) GIs. Additionally, we developed a two-step singular value decomposition (SVD)-based correction to mitigate GI-independent, biologically coherent co-variation and experimental artifacts. Finally, we employed a novel Markov chain Monte Carlo (MCMC)-based method on multiple different query screens with 30 biological replicates to rigorously quantify FP and false negative rates for qGI scores. Together, this computational pipeline enabled the mapping of a human reference GI network^62^ and provides broadly applicable tools for interpreting differential CRISPR screens.

## Results

### Genome-wide CRISPR-Cas9 GI screens in HAP1 query mutant cell lines

To map a genome-scale reference GI network (GIN) in the near-haploid human HAP1 cell line, we performed genome-wide CRISPR-Cas9 drop-out screens, measuring cell fitness as a composite phenotype following the single perturbation of every targetable protein-coding gene in a defined query mutant background (Fig. 1a). In this setting, a HAP1 query mutant cell line carrying a single loss-of-function mutation in a gene of interest is infected with a lentiviral pooled TKOv3 CRISPR library^61^, which targets nearly 18,000 protein-coding genes with four gRNAs per gene. GIs between the query gene mutation and each library gene are assessed using the qGI scoring pipeline described below. Overall, we conducted 363 genome-wide screens including 39 control screens using a wild-type (WT) HAP1 cell line, generating a dataset suitable for both accurate identification of GIs and systematic network analysis (see Methods for details)^62^. In total, this dataset comprised fitness effect measurements for approximately 23 million query gene-library gRNA double perturbations and 2.7 million single gRNA perturbations (Fig. 1b), providing the foundation for the development of the qGI score.

**Figure 1:**
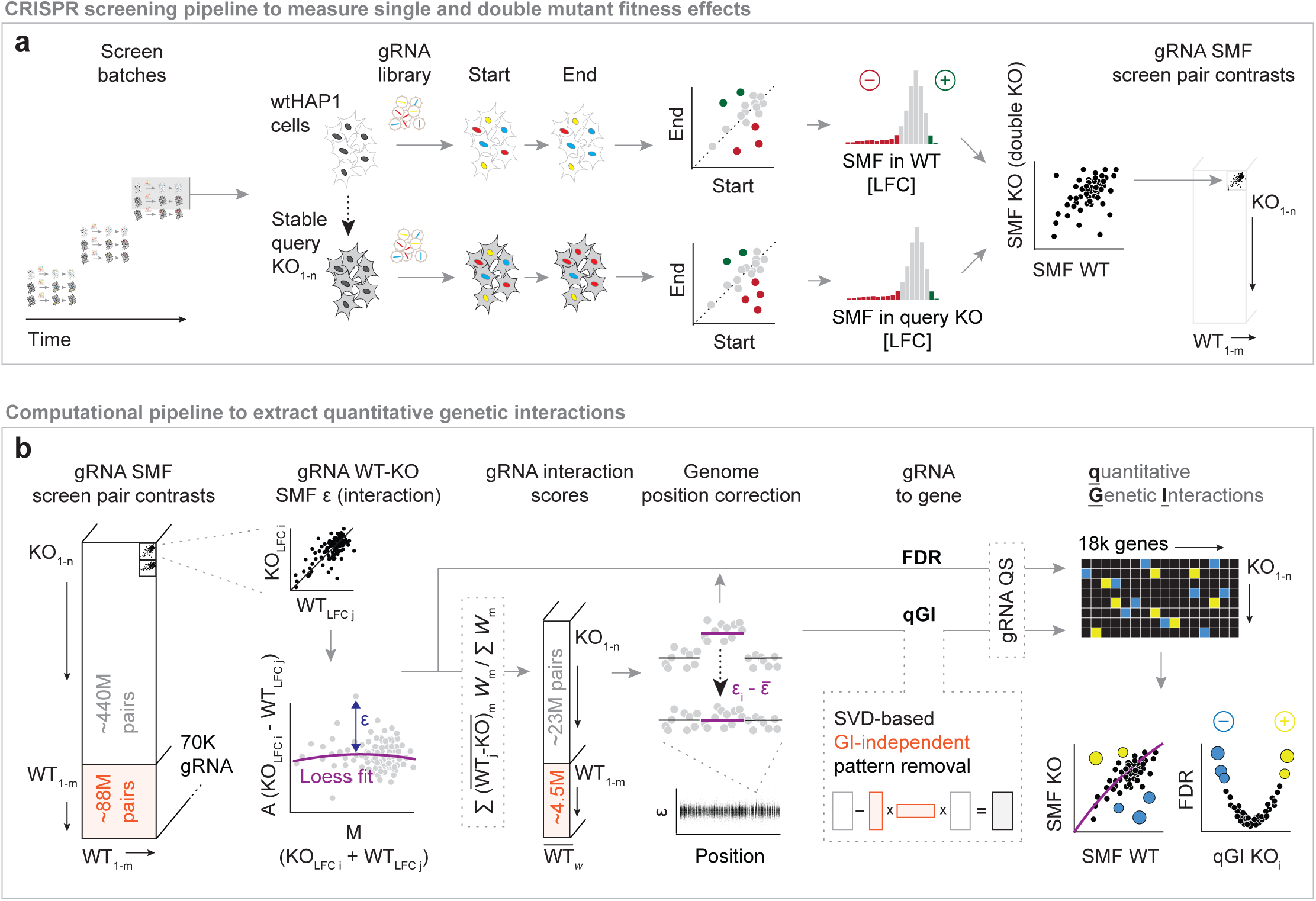
Isogenic genome-wide CRISPR-Cas9 screening to identify genetic interactions (GIs). **(a)** Quantitative GIs are identified by contrasting single mutant fitness (SMF) effects measured in HAP1 cells harboring a specific query gene knockout (KO) and control wild-type (WT) HAP1 screens as reference. Every KO-WT screen pair is compared at the gRNA level, where SMF effects are represented by the log2-foldchange (LFC) between starting and endpoint gRNA abundance, to obtain residual LFC values. **(b)** In a series of normalization steps, gRNA-level residual (ε) LFC values are processed to represent the fitness effect of the specific query KO background. A tensor of SMF contrasts of each query KO screen (n = 147 in minimal media; n = 177 in rich media) with every WT control screen (m = 21 in minimal media; m = 18 in rich media) in the same media condition provides the basis for the robust estimation of effects. Normalized residual LFC values are summarized at the gene level to compute the quantitative genetic interaction (qGI) score and an associated FDR.

### Robust identification of <u>q</u>uantitative <u>g</u>enetic interactions (qGI) from isogenic CRISPR-Cas9 screens

A GI occurs when the combined perturbation of two genes produces a phenotype that cannot be predicted from the individual effects of each gene. In our genome-wide CRISPR screens, GIs between a query gene and any library gene are reflected in the differential phenotype derived from comparing CRISPR screens performed in a query mutant cell line to those in a HAP1 WT control. However, accurate measurement of GIs, requires careful distinction between true double mutant fitness phenotypes and systematic experimental effects that introduce variation in mutant fitness phenotypes across different screens. To address this challenge, we developed the qGI score and its associated false discovery rate (FDR), which encompasses a series of normalization steps that account for artifacts in isogenic CRISPR screen data, enabling us to derive accurate, quantitative estimates of GIs. The qGI scoring methodology was applied on a large-scale, analyzing ∼4 million gene pairs and constructing a GI network encompassing ∼89,000 negative and positive GIs^62^.

The key components of the qGI scoring pipeline are: estimation of a nonlinear null model, correction of genome location-dependent artifacts, unsupervised correction of screen-to-screen variation and covariation, and estimation of the effect size and statistical significance of double mutant interactions (Fig. 1b). We provide a brief overview of these steps here. Details of the entire approach are included in Methods, and the most impactful portions of the scoring pipeline and their impact on the resulting interactions are described in more detail in the remaining sections of Results.

The qGI pipeline conceptually assumes a multiplicative null model, along with additional covariates such as single gene fitness effects. It contrasts gRNA-level fitness effects in WT control and query screens by fitting a non-linear null model to M-A-transformed fitness effects of each query-WT screen pair (see Methods for details) (Fig. 1b, Fig. 2a). Fitness effects in each screen were quantified by computing the log2 fold-change (LFC) of normalized gRNA abundance between the initial time point (T0) and endpoint (T12 to T18; 10-12 population doublings) time points, followed by basic per-screen and per-gRNA quality assurance, such as removing screens with low essential vs non-essential gene separation or gRNAs with low initial abundance (see Methods for details; Fig. S1a-k). A GI is classified as positive when the observed cell fitness exceeds the expected value and as negative when it falls below expectation, where the expectation is derived from an estimated null model. This null model substantially improved GI profile performance compared to a simple multiplicative null model as measured by the ability of profile similarity on the resulting scores to predict known functional relationships (Fig. 2b).

**Figure 2:**
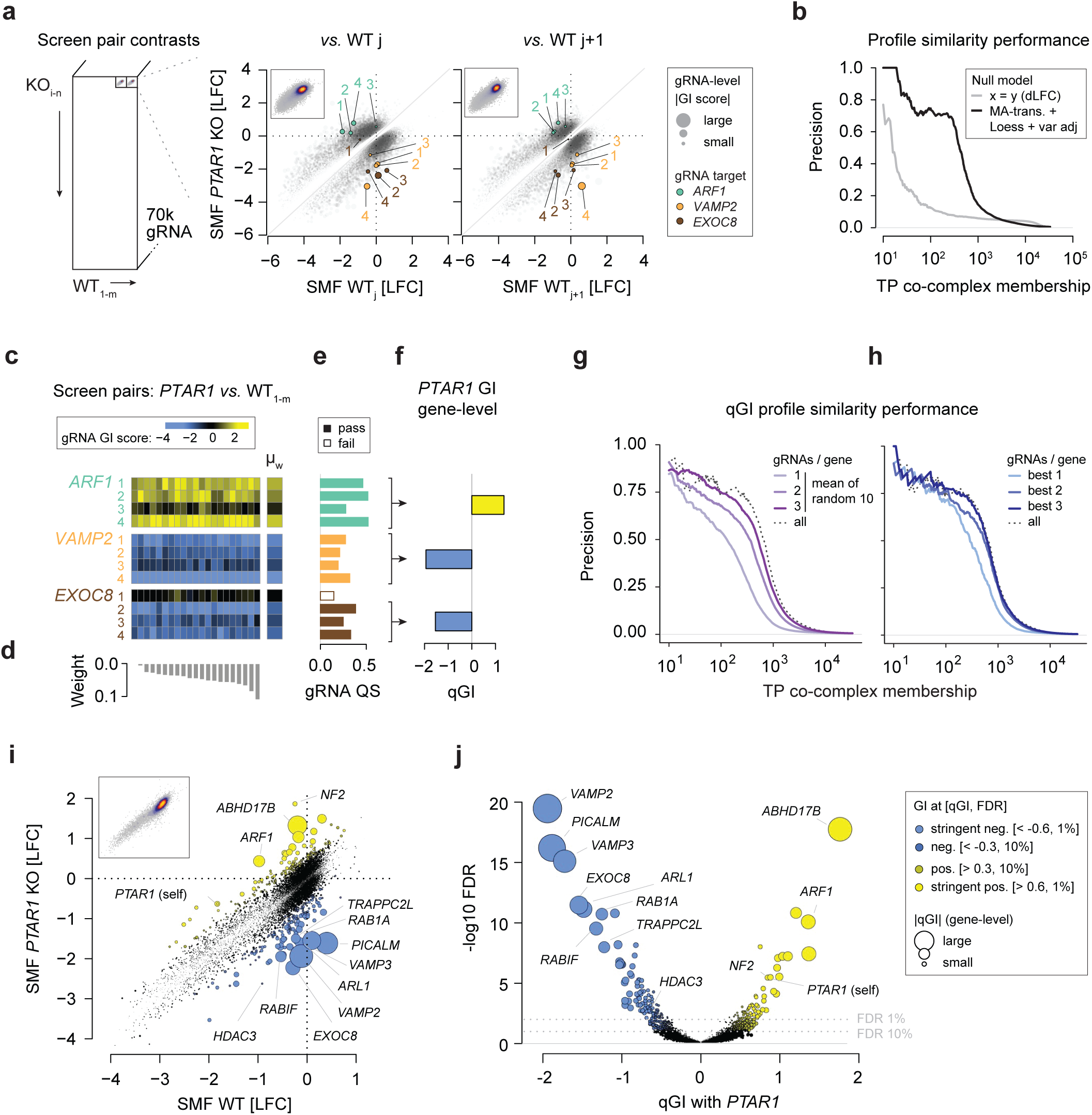
Robust identification of quantitative genetic interactions. **(a)** Pairwise comparison of gRNA-level fitness effects measured in KO query and control WT genome-wide CRISPR-Cas9 screens in HAP1 cells. Shown is an example screen performed in a *PTAR1* KO background contrasted to two WT control screens, and highlighted are the four gRNAs targeting library genes *ARF1, VAMP2* and *EXOC8*, respectively. Dot size corresponds to the fitted residual fitness effect (gRNA-level GI) between the *PTAR1* KO and each WT control screen. **(b)** Performance of dLFC (grey) and raw qGI (black) profiles across 324 query screens to identify CORUM co-complex membership. The raw qGI scores represent differential effects between query and control screen using MA-transformation, a Loess-based null model, and variance adjustment removing T0 readcount dependencies. **(c)** Complete set of residual SMF effects (gRNA qGI) between the *PTAR1* KO and all 21 minimal media reference WT control screens for the three example genes indicated. The weighted per-gRNA mean (μ_W_) was taken across the 21 KO-WT screen contrasts. **(d)** Weights of each KO-WT screen contrast are adjusted to prioritize residual effects derived from WT screens that fit the KO screen best (see Methods). **(e)** gRNA quality score (QS) for the gRNAs targeting the three example genes. The gRNA QS is derived from the sum of within-gene pairwise Pearson correlation coefficients of residual fitness effects across 324 query screens (see Methods). **(f)** Gene-level contrast of *PTAR1* KO and the mean WT control SMF effects. **(g, h)** Performance of qGI-based profiles across 324 query screens to identify CORUM co-complex membership. qGI scores are based on 1, 2 or 3 gRNAs per gene. Selected were either random gRNA targeting a specific gene (g; 10 iterations) or the gRNA per gene with the best gRNA QS (h). qGI profile Pearson correlation coefficients are used as similarity metric. **(i)** Comparison of genome-wide *PTAR1* qGI scores with SMF effects between the *PTAR1* KO and the WT control screens. Dots are scaled proportional to the absolute qGI score and colored based on the qGI score (yellow: positive GI, blue: negative GI) and FDR thresholds. **(j)** Genome-wide *PTAR1* qGI scores and the associated FDR. Dot color indicates significant genetic interactions of a library gene with *PTAR1* by considering the qGI score and the associated FDR.

To robustly quantify GIs and identify confounding patterns unrelated to the query gene that adversely impact the quality of GI estimates, we computed all pairwise differential effects between the WT screens and each query screen. This resulted in a data tensor comprising approximately half a billion measurements including more than 6,000 query-WT screen pairs, each containing around 70,000 gRNA-level differential effects (Fig. 1b). Key corrections applied by the qGI pipeline included normalization of genome location biases, SVD-based correction of GI-independent covariance patterns, both of which are described below in detail, and exclusion of inconsistent gRNAs (Fig. 1b). To quantify each query-library gene interaction, multiple query-WT screen pairs were computed, where every query screen was contrasted to all WT control screens, and gRNA-level values were summarized to estimate the effect size, the qGI score. The variance between sequence-independent gRNAs targeting the same gene and query-WT control contrasts was used to estimate the statistical significance of the interaction (see Methods for details) (Fig. 2c-f; Figure S2a, b).

Additionally, to improve the accuracy of GI scores, the qGI pipeline computes a gRNA quality score (QS) for each gene, measuring the consistency of differential effect profiles among gRNAs targeting the same gene. Using this metric, we excluded the lowest-performing gRNA for 517 genes (Fig. 2e, g, h; Fig S2c-g). We observed that a two-gRNA-per-gene library, selected based on the QS, has the potential to perform as well as the full 70,000-gRNA TKOv3 library, but four gRNAs per gene were crucial for accurate scoring of this dataset since the gRNA QS requires a large collection of screens to be computed (Fig. 2g, h; Fig. S3a-e).

Altogether, the qGI score contrasts fitness effects from genome-wide CRISPR-Cas9 query and WT control screens and applies multiple corrections to isolate the differential effects attributable to the query mutation (Fig. 2i, j). Full details of the entire qGI pipeline are described in Methods. In the following sections, we highlight two particularly important correction modules: (1) the correction of genome location-dependent artifacts, and (2) unsupervised correction of screen-to-screen variation and covariation.

### Genomic linkage of genetic interactions in isogenic cell lines

Abundant genome position-linked false positives (FP) in CRISPR screens have previously been described and can arise through multiple mechanisms^54,63^. One prominent example is gene function-independent CRISPR-Cas9-induced DNA repair-associated toxicity, which increases with DNA copy number (CN) and can produce cell line-specific fitness effects and introduce genomic linkage-driven coessentiality among neighboring genes^50,53,54,64^. Moreover, CRISPR-Cas9-induced DNA double-strand break-mediated loss of chromosome ends can generate a collateral knockout of essential genes^63^. While our isogenic screening platform is designed to avoid such artifacts, we specifically tested if our GI screens in HAP1 cells showed evidence of genome position-linked effects. Specifically, we tested the impact of two scenarios on our ability to identify GIs: (1) ploidy changes where all cells in the pool of the haploid HAP1 cell line spontaneously diploidized and (2) ploidy differences restricted to defined genomic regions in the query mutant and HAP1 WT control screens.

Sometimes, haploid HAP1 cells (1n) can undergo diploidization (2n) during query clone generation or screening. Given the CN–dependent toxicity of CRISPR-induced DNA damage, contrasting the CRISPR screens performed in HAP1 WT cells that remained haploid with those that became diploid during the experiment, or those that were diploid from the outset, may introduce differential artifacts. Such effects would likely cause FP GIs when contrasting query mutant with HAP1 WT screens that also display different ploidy status. However, neither the number of gRNAs producing strong fitness defects nor the number of detected differential fitness effects differed between the three ploidy classes (Fig. S4a-d). Thus, whole-genome ploidy differences at the start or end of a screen are unlikely to systematically generate FP GIs using our platform.

Next, we examined a 30 Mbp region on chr15 that is heterozygous diploid in most HAP1 query mutant clones as well as in the HAP1 WT control cells but has been engineered to be haploid in a few so-called eHAP1-derived query mutant clones^65^ (Fig. 3a). Notably, for the *WDR73* query gene cell line, which is haploid on the entire chr15, we found that gRNAs targeting 222 genes within this region exhibited markedly positive qGI scores, on the order of a ∼30-fold increase above the global background density (Fig. 3b) (hypergeometric test; p < 2.2 x 10^-16^) before correction (Fig. S5a-c). This observation suggests that unequal local CN differences between a query and WT control screen pair can cause high rates of FP GIs.

**Figure 3:**
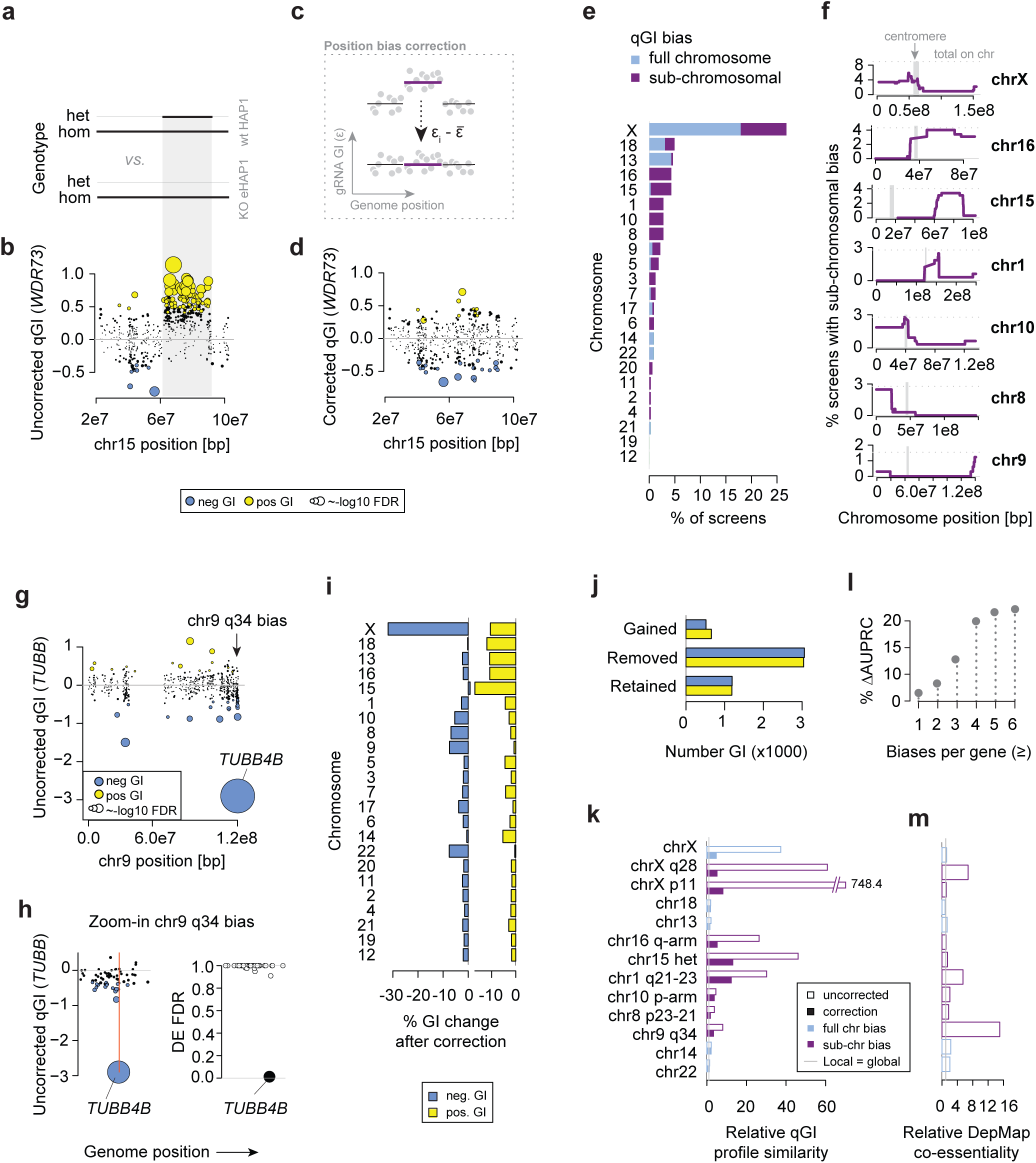
Correction of genome position-linked false positive genetic interactions. **(a)** Homozygous haploid and heterozygous diploid genotype of chr15 in eHAP1-derived *WDR73* query clone and HAP1 WT control cells. **(b)** Uncorrected WDR73 qGI scores on chr15. Dot size corresponds to -log10 FDR of the qGI score, and colors indicate significant GIs (|qGI| > 0.3, FDR < 0.1). **(c)** Schematic illustration of the correction of genome position-linked biases through mean-centering of gRNA-level interaction scores. **(d)** Corrected *WDR73* qGI scores on chr15. Dot size corresponds to -log10 FDR of the qGI score, and colors indicate significant GIs (|qGI| > 0.3, FDR < 0.1). **(e)** Percentage of HAP1 query gene KO screens with a residual (ε) SMF (uncorrected qGI) biases on each chromosome (purple) or a sub-chromosomal region (green). 100% corresponds to 324 screens. **(f)** Frequency of affected genomic regions on the seven chromosomes with the most sub-chromosomal biases. The upper dotted line indicates the frequency of total sub-chromosomal biases on that chromosome. **(g)** Uncorrected *TUBB* qGI scores on chr9. Dot size corresponds to - log10 FDR of the qGI score, and colors indicate significant GIs (|qGI| > 0.3, FDR < 0.1). **(h)** Uncorrected *TUBB* qGI scores and differential expression (DE) in the affected 64-gene chr9 q34 region. Blue (qGI; left) and black (DE; right) mark significant scores. Significant DE is called at FDR < 5%. Orange line marks genome location of *TUBB4B*. **(i)** Net percentage of GIs lost and gained after correction for local biases in each chromosome across all 324 screens. **(j)** Number of positive (yellow) and negative (blue) GIs (|qGI| > 0.3, FDR < 0.1) across all biased regions in the 324 screens that were gained, removed and retained after correction. **(k)** Relative local density of qGI profile similarity of genes in the genomic region with the most chromosomal and sub-chromosomal biases before and after correction. A value above 1.0 indicates that strong (z > 4) profile similarities are observed more often between genes encoded in an affected genomic region than between genes not encoded in the same region. **(l)** Change of the area under the PR curve when GO BP relationships are predicted through qGI profile correlation. The analyses were focused on all library genes affected by at least a certain number (1-6) of called biases. **(m)** Relative local density of highly similar (z > 4) co-essentiality profiles across 1095 different cell lines (DepMap 23Q2 release) with the affected regions in isogenic HAP1 screens.

We developed an unsupervised algorithm to systematically detect and correct qGI scores associated with genomic regions showing systematic biases toward positive or negative GI scores (see Methods). The qGI scoring pipeline mitigates these biases by mean-centering gRNA-level scores within these localized regions (Fig. 3c). For instance, correcting the GI scores in the 30 Mbp region on chr15 in the *WDR73* KO screen retained only 7 of the 77 positive interactions (qGI > 0.3; FDR = 10%) while simultaneously identifying 11 new negative interactions (qGI < -0.3; FDR = 10%), none of which passed the threshold without this correction (Fig. 3d).

When applied to all query screens and all chromosomes, the algorithm detected genome position-linked biases in uncorrected qGI scores affecting 75,739 gene pairs (1.3% of all pairs tested), involving 42% of all query gene screens and 56% of all library genes. We grouped these biases into two categories: biases that affected the full chromosome or biases that affected only subchromosomal regions. The frequency of these biases varied by chromosome, with chrX most frequently affected (26.9% of query clones) (Fig. 3e). Some chromosomes exhibited a preference for either full-chromosome or subchromosomal biases (Fig. 3e; Fig. S5d). For example, chr13, chr18, and chrX predominantly displayed whole-chromosome biases, while other chromosomes exhibited mostly subchromosomal effects (Fig. 3f; Fig. S5e, f). When subchromosomal biases occurred, they clustered in specific regions, suggesting that those regions are prone to genetic or epigenetic instability in HAP1 cells (Fig. 3f). Some subchromosomal biases, such as those on chr16 or chr10, affected one but not the other chromosomal arm (Fig. 3f). On chrX, which otherwise predominantly showed whole-chromosome biases, subchromosomal biases frequently occurred at p11 and q28 (Fig. 3f).

We also observed rare, genomically linked biases in individual cell lines, such as a 64-gene region at the end of chr9 that encodes *TUBB4B* in the *TUBB* query gene clone (Fig. 3f, g). In HAP1 WT cells, *TUBB4B* is highly expressed but not essential when depleted. In the *TUBB* clone, *TUBB4B* expression is upregulated (Fig. 3h), which presumably compensates for the loss of *TUBB*, and *TUBB4B* becomes highly essential, scoring as a strong negative GI (Fig. 3g, h). This buffering relationship was further supported by data from the CCLE cell line panel (Fig. S5g-i) and in a Perturb-Seq experiment in K562 cells^66^. Besides *TUBB4B*, 16 neighboring genes in this region exhibited negative GIs, a ∼25-fold enrichment over background. Since these genes did not show altered mRNA expression in the *TUBB* clone nor dependency on *TUBB* mRNA level in the Cancer Dependency Map (Fig. 3h, Fig. S5j), we suspected these neighboring genes were unlikely to contribute to buffering of *TUBB*. We found that 15 of these 16 genes were located to the centromere-side of *TUBB4B* (Fig. 3h), which suggests that their observed fitness defects may be due to occasional DNA double-strand break-mediated loss of the end of chr9, following a mechanism previously described by Lazar and colleagues^63^. Such a loss of the end of chr9 would likely knock out *TUBB4B* collaterally and produce many FP negative GIs.

Since this putative chr9 mechanism differs from what we observed in the heterozygous diploid region of chr15, where we observed significant mRNA deregulation in the biased region, we performed mRNA sequencing analyses in 42 of our HAP1 query mutation clones covering 77 biased regions. Only about 20% of these regions showed differential mRNA expression in the affected region and, of those, only biases on chr15 and chr8 (Fig. 3f) consistently showed differential expression, which is coherent with a CN hypothesis. Notably, only 1 of 26 tested biases on chrX showed differential expression. Together, these findings suggests that the observed genome position-linked biases may only be attributed to local CN differences between query and control screens in relatively few cases.

The unsupervised correction of these genome position-linked biases implemented in the qGI scoring pipeline affected each chromosome differently by removing different fractions of negative and positive GIs (Fig. 3i, k). For instance, about 30% of all negative and 10% of all positive GIs involving genes on chrX were removed following correction (Fig. 3i). Overall, the correction removed ∼6,000 FP GIs (∼3%), which exceeded the number of true positive GIs in the biased regions by approximately 3-fold (Fig. 3j; Fig. S5l).

We also evaluated the effect of genome position-linked GIs and their correction as implemented in the qGI scoring pipeline. The set of GIs for a given library gene provides a phenotypic signature indicative of gene function, connecting genes that share similar GI profiles. We observed that genes co-localized in genomic regions shared many uncorrected GIs, which results in inflated GI profile correlations and has the potential to confound functional analyses (Fig. 3k) (see Methods). For instance, the 222 genes in the duplicated region of chr15 region in HAP1 cells were nearly 50 times more likely to have highly correlated GI profiles. This was reduced after correction, which also substantially improved the predictive performance of these GI profiles (Fig. 3k, l). Indicative of a more general issue in cultured human cancer cell lines, we noticed that several regions with detected biases in HAP1 also had strongly increased relative local co-essentiality density in the Cancer Dependency Map, despite sensitive corrections for proximity bias included in the current DepMap scoring pipelines^53^ (Fig. 3m).

In conclusion, we identified regions in the HAP1 genome prone to GI score biases that can generate numerous putative FP GIs and inflated profile similarities when uncorrected in our HAP1 CRISPR screens. The qGI position-correction module can detect and normalize these biases. A subset of these effects can be explained by chromosomal abnormalities and/or collateral chromosome arm loss, but our analysis suggests other, still uncharacterized, mechanisms are responsible in a number of cases.

### Genetic interaction-independent patterns in genetic interaction screens

Previous work on yeast GIs demonstrated that technical artifacts can generate non-random patterns that can explain substantially more variation than the GIs we hope to score^13^. Given that there have been insufficient previous data capturing large-scale GI networks from isogenic CRISPR-Cas9 query screens to comprehensively characterize technical artifacts, we generated a large collection of WT control screens. These screens were generated with two potential types of artifacts in mind (Fig. 4a): (1) between-screen variation due to differences in query mutant growth dynamics, and (2) between-screen variation due to systemic experimental and random differences. Regarding growth dynamics, one of the major factors that varies in human cell culture is the growth rate of a particular HAP1 query mutant. Growth rate can impact comparison of CRISPR screens substantially^60,67^. The number of doublings of a query mutant cell population is tracked in our screening protocol, and screens were conducted following a strictly defined protocol, controlling as many factors as practically possible. But due to the multiple serial passages, precise control of this factor across hundreds of screens is not practically feasible. To understand this variation, in addition to single endpoints, we collected multiple intermediate timepoints from several control screens, sampled every 3-4 days, simulating those that were unintentionally either undergrown (fewer cell doublings than average) or overgrown relative to the average control. Regarding general between-screen variation (2), there are inherently uncontrollable elements that may drift over the course of a project (e.g., staff conducting the screens, virus batch, plasmid batch, media batches) which can introduce both random and systematic variation. To model this variation, HAP1 WT control screens were periodically completed using the same screening protocol over several years of screening.

**Figure 4:**
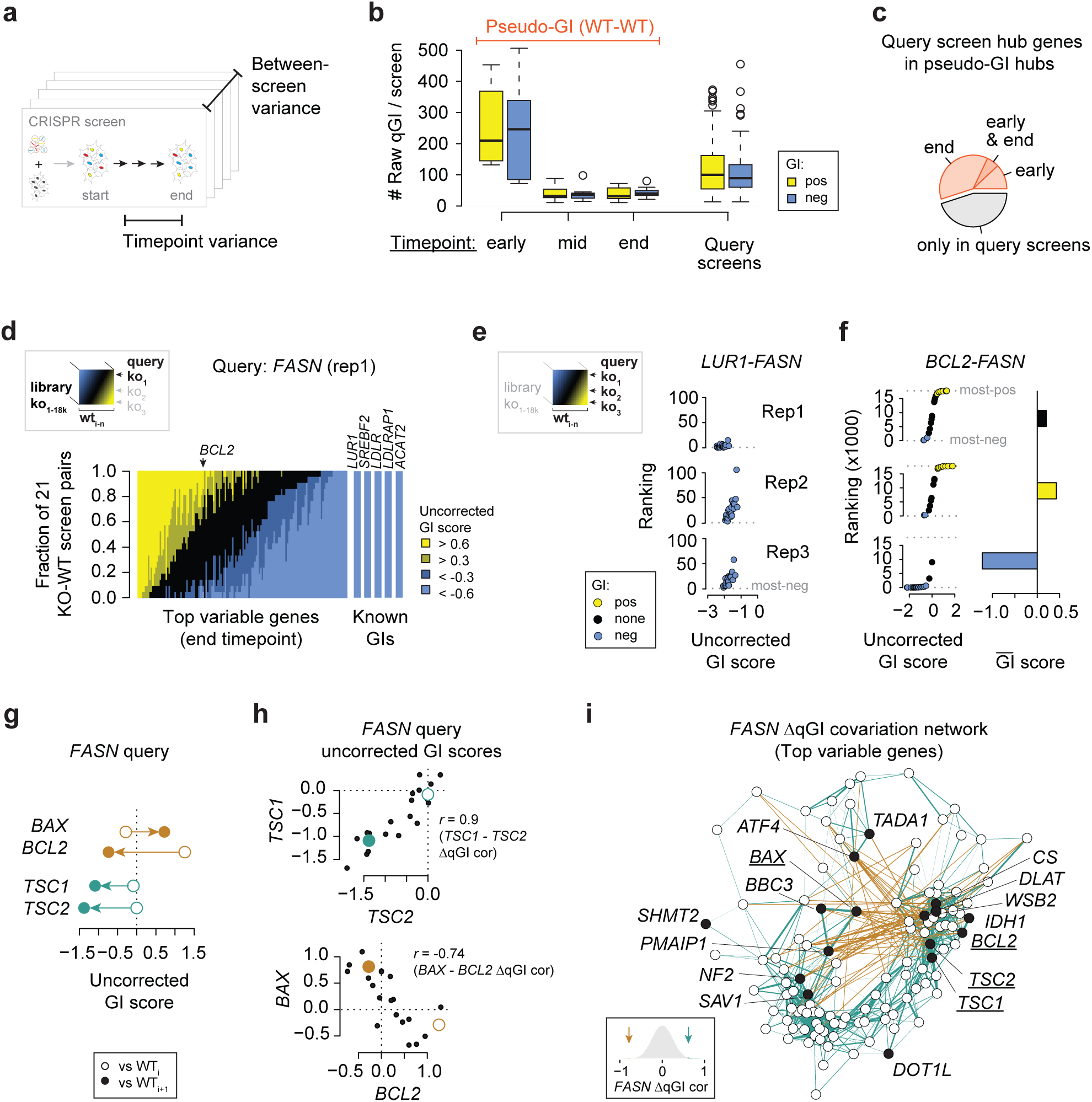
Genetic interaction-independent patterns in genetic interaction screens. **(a)** Schematic illustration of sources of library gene-specific variance. **(b)** Number of GIs for early timepoint (T6 – T10), mid (T11 – T17) and end point (T18, T19) control simulations and query screens. The qGI are uncorrected and a GI is counted for |qGI| > 0.6. Simulations iteratively contrast a held-out control screen against the set of all other control screens (see Methods). **(c)** Hub GI genes overall between simulated “pseudo-GIs” and query screens. In each data set, the top 100 genes with the highest absolute GI degree (|qGI| > 0.6) are tested for overlap. **(d)** Fraction of positive and negative uncorrected qGI scores of ∼140 highly variable genes and 5 high confidence negative GIs with *FASN* when in all 21 KO-WT contrasts. The highly variable genes were selected because they exhibited the highest fitness effect-adjusted variance between control screens or exhibited high covariance in the control screens. **(e, f)** Ranking and qGI scores (uncorrected) between *FASN* and *LUR1/SPRING1* (known interactor) or the highly variable gene *BCL2* in the 21 WT contrasts with three independent FASN KO screens. The GI spectrum thumbnail indicates that highly variable genes such BCL2 vary in their qGI score depending both on the WT control as well as on the query KO (here *FASN*) screen. Per-*FASN* replicate mean *BCL2* qGI scores are illustrated in the barcharts (f). **(g)** Uncorrected qGI scores between four highly variable genes and *FASN*. Shown are scores from the first *FASN* KO screen replicate contrasted to two different WT control screens. **(h)** Uncorrected qGI scores between the *FASN* query gene and *TSC1* and *TSC2* (top) and *BCL2* and *BAX* (bottom) when considering all 21 WT control screens. The scores shown in (g) are highlighted. Pearson correlation coefficients indicate how raw qGI scores of two genes co-vary across the 21 WT reference contrasts. **(i)** Covariation network illustrating positive (green) and negative (brown) correlation between the highly variable genes for the residual qGI scores between the replicate 1 *FASN* KO screen and the 21 WT control screens. Edges are shown for Pearson correlations above 0.6 or below -0.75. The histogram shows the genome-wide distribution of the *FASN*-specific residual qGI covariation network.

With this relatively large collection of control screens, we performed a simulation to characterize random and systematic variation associated with our GI screening platforms. Specifically, we successively held out one of the control screens and used this “decoy” screen as a test query screen to produce “pseudo-GI” scores, i.e. differential effects that resulted from contrasting that decoy to the average of all other control screens. For illustration, we focus on the media condition with the larger number of control screens (n = 21; selected from 39 total control screens). This scoring process included all of the components of the qGI score except for the SVD-based GI-independent pattern removal components (Fig. 1b). We then analyzed pseudo-GI scores for random or systematic variation and used this information to quantify the expected number of FP GIs.

We first focused on intermediate sampling points to characterize the residual effects produced by differences in the query growth dynamics and/or screen sampling time. We found that early time points yielded a high density of pseudo-GIs, which even surpassed the average query screen GI density (Fig. 4b). Endpoint samples from this control set, which model screen-to-screen variation effects unrelated to sampling time effects, displayed fewer pseudo-GIs, but still resulted in a substantial number of FP GIs (31 and 44% of the median positive and negative interaction degree, respectively) relative to the average number of GIs observed across query mutant screens (Fig. 4b). A number of the library genes classified as genetic network hub genes, which showed many true GIs, showed a relatively large number pseudo-GIs in control screen simulations (Fig. 4c). This effect was greater for screen-to-screen variation, suggesting that uncorrected between-screen systematic variation can lead to FP GIs and imprecise control of the sampling time for a particular query cell line may only contribute a small fraction of those FP GIs.

To test if between-screen variation effects between endpoint samples can produce FP GIs at the magnitude of strong true GIs, we contrasted each query mutant screen with each of the WT screens (n=21 in minimal media), which resulted in 21 differential effect measurements for each query gene. For illustration, we focused on two query genes with well-characterized functions, *PTAR1*, which participates in vesicle trafficking, and *FASN*, which codes for Fatty Acid Synthase and assembles long-chain fatty acids. Specifically, we wanted to understand whether genes that showed positive or negative qGI scores in one query-control contrast also did so in the remaining 20 contrasts for the same query gene. Strong negative and functionally coherent GIs with *PTAR1* or *FASN* were found in every one of the 21 contrasts, suggesting that a simple query-control screen comparison can robustly identify true interactions (Fig. 4d, Fig. S6a). For instance, the true negative GI between *PTAR1* and a vesicle trafficking regulator *VAMP2* showed a strong negative qGI score in each of the 21 contrasts, ranking among the top most negative interactions in each case (Fig. S6b). In the case of the *FASN* query, we examined three replicate screens and followed the GI between *FASN* and *LUR1*^32^, which was among the most negative GIs in each of the 63 contrasts (Fig. 4e). Importantly, some of the highly variable library genes across these contrasts resulted in scores that were similar in magnitude in specific contrasts to reproducible interactions but even switched the sign of the GI between contrasts (Figure 4d, Figure S6a). For instance, the interactions between *PTAR1* and the highly variable gene *BCL2* showed strong negative or strong positive interactions ranging between qGI = -1.33 (15^th^ most negative interaction in the genome) to qGI = 0.84 (53^rd^ most positive) (Figure S6c).

To explore if our collection of control screens was sufficient to mitigate FP GIs of highly variable genes, we mean-summarized the 21 KO-WT contrasts for each of the three biologically independent *FASN* query screens, which resulted in a single control profile robust against control screen variation. The three resulting *BCL2-FASN* GI scores ranged from -1.2 (very strong negative GI) to 0.4 (moderate positive GI) (Fig. 4f). This showed that highly variable genes across WT control screens also vary frequently in different query mutant screens, suggesting that between-query screen variation requires additional correction to mitigate FP GIs.

We next asked whether the variation observed in control screens was independent across library genes, or instead, showed *covariation* between library genes. To examine for potential covariation, each query gene screen was individually contrasted against the 21 WT control screens, resulting in 21 differential effect measurements for each query x library gene combination. Surprisingly, we found strong covariance between biologically coherent groups of genes with high between-screen variation. For instance, the anti-apoptotic regulator *BCL2* and the pro-apoptotic regulator *BAX*, which can form a dimer with *BCL2* in response to stress to trigger apoptosis, showed the expected opposing qGI score across many *FASN*-WT contrasts, but the GI sign for *BCL2* and *BAX* flipped simultaneously in some *FASN*-WT contrasts (Fig. 4g). In the 21 *FASN*-WT contrasts, the *BCL2-BAX* qGI scores were strongly anti-correlated (Fig. 4h). In another example, *TSC1* and *TSC2*, which form a dimer to inhibit nutrient-dependent growth regulation through mTORC1, both interact negatively with *FASN* in one WT contrast, but a GI is absent in another WT contrast, which represents a residual GI pattern with a strong positive correlation (Fig. 4g, h). We computed correlations across the entire set of highly variable genes to construct a global covariation network for the *FASN* query (Fig. 4i). Overall, this network was highly structured and contained several functionally coherent modules, including those that are known to be central to cell growth and cell death phenotypes (Fig. 4i). These findings suggest there is substantial systematic variation and covariation present in genome-wide CRISPR screens in biologically replicated conditions whose effect size is equal or greater in magnitude than the interaction effects being measured. Furthermore, this covariation reflects biologically coherent modules. Thus, identifying true GIs (i.e., differential effects that are specifically caused by the introduced query mutations) requires careful normalization of this systematic variation.

### Correcting for genetic interaction-independent fitness patterns

To correct for the variation and covariation observed across control screens, we developed a two-step singular value decomposition (SVD)-based normalization module for the qGI pipeline that selectively removes covariation patterns resulting from systematic variation in query screens. First, this module removes pseudo-GI signal caused by library genes with variable single mutant fitness identified in WT GI simulation analysis. A matrix containing pseudo-GI scores derived from the WT simulation analysis, described above (Fig 4), was decomposed using SVD (Fig. 5a) (see Methods), and the patterns corresponding to the resulting singular vectors were removed from our qGI data matrix. Specifically, we computed the projection of the query screen raw GI scores onto the pseudo-GI-derived singular vectors and subtracted the result from the raw GI scores to produce a normalized qGI score (Fig. 5a) (see Methods for details). To illustrate the effect of this correction, we focused on library genes shown to have the most variable single mutant fitness effects across WT control screens (Fig. 4d, i) and arranged them into a similarity network based on their covariance structure (Fig. 4i). We visualized the correction of individual query screen interaction profiles on this similarity network by plotting the ΔqGI score on this network, the difference between normalized and unnormalized qGI scores. As described above, this similarity network groups genes into coherent functional modules. For each query screen, corrections showed coherent patterns on this similarity network but varied substantially in strength and magnitude from screen to screen (Fig. 5b, c). For example, GIs between *FASN* and genes in the cluster *INPP4A, INPPL1, FBXW7, KCTD5* and *HDAC3,* or *TSC1* and *TSC2* were corrected to become less negative (Fig. 5b). In contrast, the cluster *TADA2B, TADA1, HK2, SLC16A1* and *ATF4* was strongly corrected to show less positive GIs (Fig. 5b). Other query screens showed distinctive correction patterns relative to *FASN*, each with a different combination of negatively and positively corrected library genes (Fig. 5c). This screen-based correction is driven by the covariance observed across the controls, but which collection of library genes is corrected in any given screen can vary substantially, suggesting different GI-independent patterns emerge with varying strengths in each query screen.

**Figure 5:**
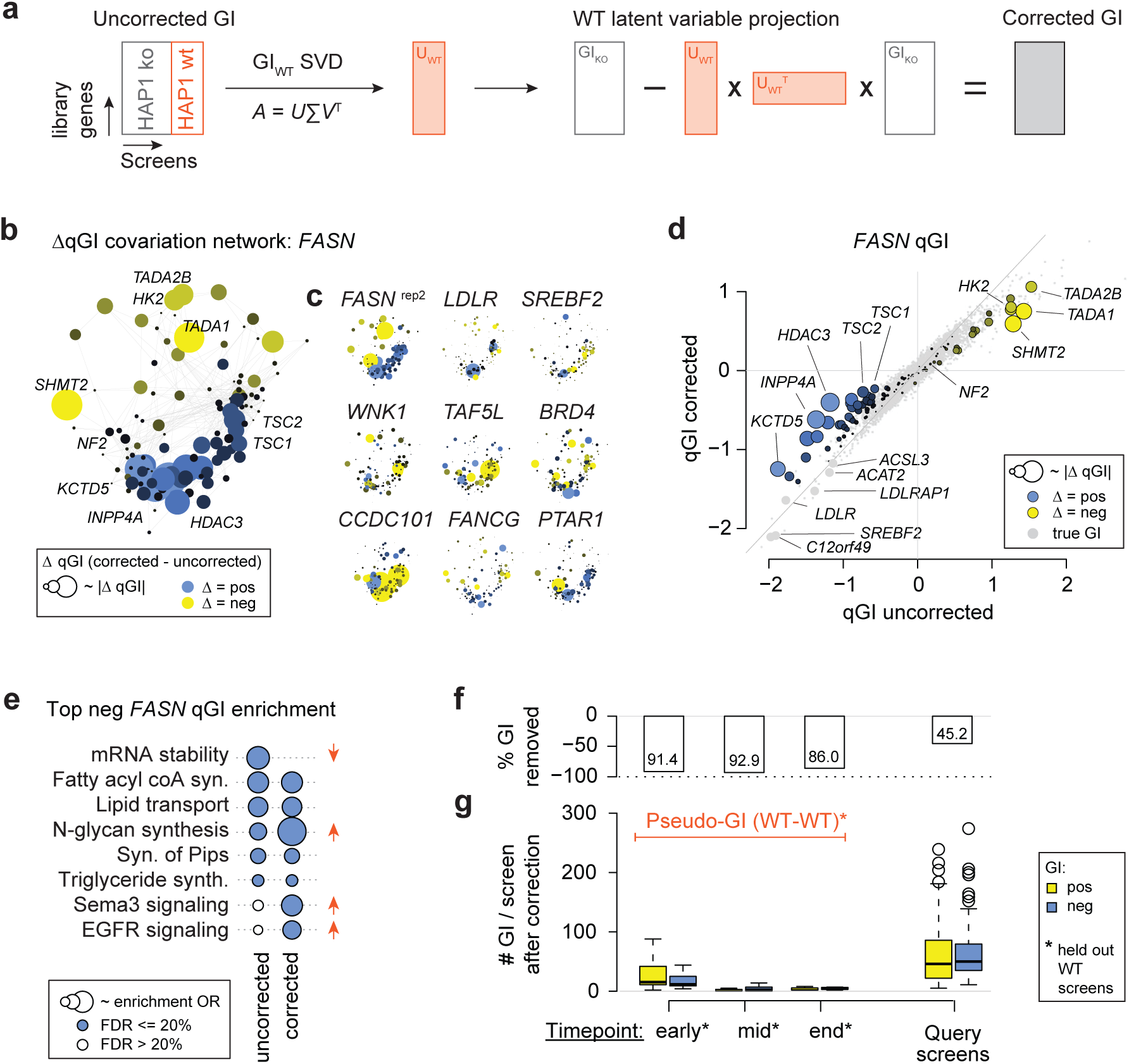
Correcting genetic interaction-independent patterns in genetic interaction screens. **(a)** Schematic illustration of the SVD-based removal of WT-derived variance on query screens: for each of the ∼70k gRNAs, latent variables describing GI-independent variation are derived from the WT control screens and projected onto and removed from raw qGI scores measured on query screens. **(b)** *FASN*-specific covariation network for the highly variable genes (see Fig. 4i). Genes are connected by their residual qGI profile correlation across the 21 *FASN*-WT contrasts. Dot size corresponds to the gene-specific change of the qGI score. A correction towards a more positive value (less negative qGI score) is shown in blue, and correction toward more negative (less positive qGI score) in yellow. **(c)** Correction thumbnail laid out as in (b) for 9 different query screens. The gene-specific change of the qGI score represents the change in a given query screen. **(d)** *FASN* qGI scores before and after correction for GI-independent patterns. Shown are 17,804 genes, and the highly variable genes are colored and a subset are labeled. Dot size corresponds to the gene-specific change of the qGI score. A correction towards a more positive value (less negative qGI score) is shown in blue, and correction toward more negative (less positive qGI score) in yellow. **(e)** REACTOME pathway enrichment of the top 100 negative and positive qGI scores in the *FASN* query screen before and after correction. All pathways enriched at an FDR of 20% before or after are shown. **(f)** Total net change in the number of GIs after correction (Fig. 4b vs Fig. 5g) **(g)** Number of GIs for early timepoint (T6 – T10), mid (T11 – T17) and end point (T18, T19) control simulations and query screens after correction (compare to Fig. 4b). The qGI are uncorrected and a GI is counted for |qGI| > 0.6. Simulations iteratively contrast a held-out control screen against the set of all other control screens.

A major determinant of how strongly a library gene is corrected is the combination of their between-screen variance, covariation patterns, and the query-specific interaction effect. Notably, this allows this correction to be both sensitive and specific. For instance, while scores of highly variable genes were substantially modified, functionally coherent negative GIs with *FASN*, such as with lipid transport or lipid biosynthesis genes *LDLR, LDLRAP1, SREBF2, LUR1/SPRING1*, were not affected by this correction (Fig. 5d). This was also reflected by functional enrichment among the top negative GIs with *FASN*, which showed removal of GIs with genes involved in mRNA decay but retained GIs with genes in lipid metabolism-related pathways (Fig. 5e). The same trend was observed in the *PTAR1* query screen, where highly variable genes showed large corrections while functionally coherent negative GIs with the vesicle trafficking genes *VAMP2, VAMP3, EXOC8,* or *ARL1* remained unchanged (Fig. S6e). Interestingly, for the *PTAR1* query, the correction removed a significant enrichment for mTOR signaling among negative interactions with *PTAR1* while retaining enrichments for Golgi vesicle biogenesis, which reflects *PTAR1* function in vesicle trafficking (Fig. S6f). Overall, the first SVD-based correction module removed the majority (∼90%) of pseudo-GIs, reflecting time sampling and screen-to-screen variation observed in our simulation experiment (Fig. 5f, g). Overall, a surprising 45.2% of GIs in query screens were removed, indicating a pervasive false positive signal in uncorrected GI scores (Fig. 5f, g). Among the GIs that were removed, positive GIs were more frequent (∼57%), but the proportion of positive and negative GIs removed varied substantially between different library genes. For example, some genes, including *HDAC3, MORC2* and *NF2*, primarily lost negative GIs, while others, such as *WSB2* or *PTDSS1*, lost more positive GIs.

The second normalization step involves removal of the most dominant signal from the query x library gene matrix in an unsupervised manner (Fig. 5a). This step was designed to remove additional systematic experimental effects in the GI data. Given that GIs are rare, any low-dimensional pattern that explains a substantial portion of variance in the data likely represents technical artifacts, not true GIs. Previous studies large-scale GI studies demonstrated that normalization of such patterns (e.g., the strongest principal components) from interaction data can improve the quality and specificity^13,53,68^. To test how the strongest principal components contribute false positive signal in our context, we measured how well GI profiles reconstructed co-membership of gene pairs in orthogonal functional standards. We found that removing the first 4 principal components from the data, which account for ∼11% of the total variance (Fig. S6g), increased the performance of qGI scores to detect co-complex and co-pathway membership (Fig. S6h, i). In this approach, the top SV-removal correction, removed positive and negative GIs in a relatively balanced fashion. However, the fraction of positive to negative GIs was highly dependent on the library gene. Notably, this module removed many positive GIs involving components of the respiratory chain complex I, such as *NDUFAF6, NDUFA6* or *NDUFB8*, which we have shown previously to dominate CRISPR screen data in part due to technical artifacts^67^. In contrast, a higher number of negative GIs involving SWI/SNF complex components, including *SMARCC1, SMARCB1* or *SS18*, were removed.

In addition to our SVD-based normalization approach, which was integrated into the qGI scoring pipeline, based on the analysis of variation and covariation across the control screens, we specifically identified a set of library genes associated with highly variable single mutant fitness phenotypes. We defined two sets of genes: the first a “core” set of 80 highly variable fitness genes with the highest variability, and a second “expanded” set of 426 library genes with more subtle effects (see Data File S2 in^62^). While our proposed normalization approach can moderate the effects of systematic variation, we suggest that caution should be used in interpreting single or double mutant phenotypes for these genes, particularly for the “core” variable fitness set.

### Cumulative impact of qGI normalization on interactions

As described above, we characterized several biological and technical artifacts arising from our genome-wide CRISPR/Cas9 screening platform that obscure true GIs and developed a collection of normalization procedures, which collectively form the qGI pipeline. In particular, we evaluated three variants of the qGI method relative to the complete approach, each with progressively more limited functionality: one without the removal of the top four SVs, one without any SVD-based correction, and one without the non-linear null model based on M-A-transformed fitness effect data (referred to as simple dLFC). Each of those ablation experiments suggested that the qGI score correction modules significantly improved the functional information derived from the resulting qGI scores, and cumulatively, they had a major impact on the qGI profiles’ capacity to predict co-complex, co-pathway and co-GO BP memberships (Fig. 6a, Fig. S7a, b). In addition, each correction module reduced the number of strong GIs overall by more than 80% (Fig. 6b). This contradicts the frequent assumption that technical artifacts in CRISPR screen data mostly affect moderate and weak phenotypes and that strong effects reflect the true signal of interest. In total, proportionally more positive than negative GIs were removed (Fig. 6b), the positive-to-negative GI ratio of each screen resolved, becoming more balanced upon normalization (Fig. 6c). For example, for the *WNK1* query screen, a simple dLFC measurement generated large numbers of negative GIs due to a non-linear relation to the WT control reference set (Fig. 6d). Applying an accurate null model as well as the additional qGI corrections removed many strong effects while retaining strong, functionally coherent GIs, such as those expected between the *WNK1* paralogs *WNK2* and *WNK3*. This trend for the *WNK1* query, where a large fraction of genes with a specific fitness effect are called as GIs when no corrections are used due to the non-linear query-control relation of genome-wide fitness effects was broadly representative of corrections for other queries as measured via a global correlation analysis (Fig. 6e).

**Figure 6:**
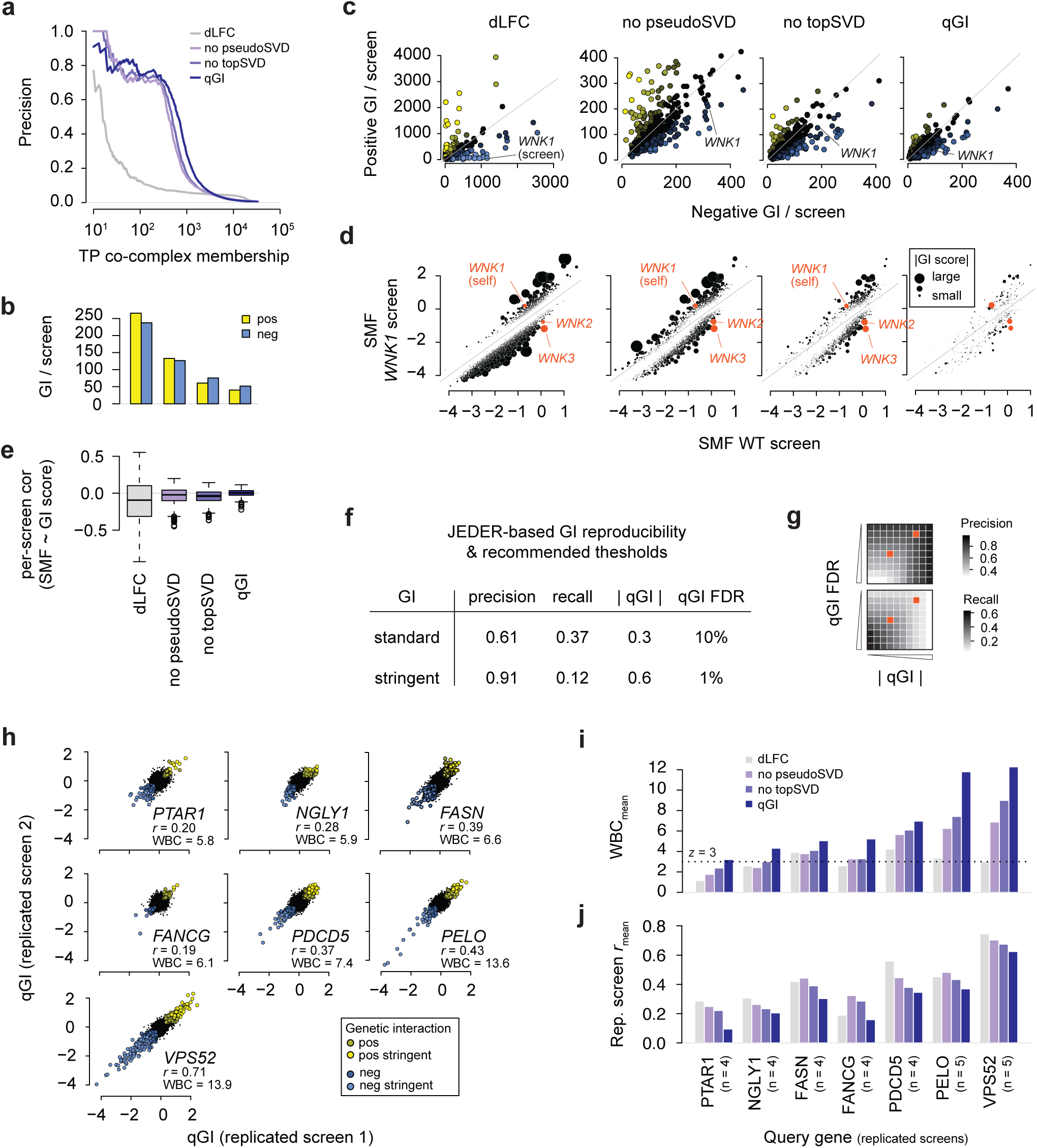
Cumulative impact of qGI normalization on genetic interactions and reproducibility analysis. **(a)** Increasing precision-recall performance of qGI profiles at different normalization steps of the qGI pipeline to reconstruct known CORUM co-complex memberships. Shown is the simple differential (d)LFC, a fitted raw qGI version without removal of GI-independent patterns including the top SV correction, a version without removal of top SVs, and the completely corrected qGI score. **(b)** Mean number of GIs per query screen at different stages of normalization of the qGI pipeline. A GI is called at |qGI| > 0.6. **(c)** Per-query screen comparison of the number of positive (qGI > 0.6) or negative (qGI < -0.6) GIs at different normalization stages of the qGI pipeline. Blue indicates a higher ratio of negative, yellow a higher ratio of positive GIs. **(d)** Differential fitness effects of the library genes in the *WNK1* query screen at different normalization of the qGI pipeline. Fitness effects in WT HAP1 control are plotted against fitness effects in the query screen. Dot size is scaled to the absolute interaction score. The paralogs of *WNK1* are labeled. **(e)** Per-query screen correlation of the fitness effect and interaction scores at different normalization stages of the qGI pipeline. **(f)** Selected thresholds for the GI effect size, the qGI score, and significance, the qGI-associated FDR, for standard and stringent GIs. The JEDER-derived precision and recall are highlighted in (g). **(g)** Precision and recall of positive and negative GIs of the 7 sets of replicated query screens (see i, j) at different qGI and FDR thresholds. The precision was estimated using a standard-free MCMC-based method to determine FP and FN rates. The set of qGI and FDR thresholds for standard (|qGI| > 0.3, FDR < 10%) and stringent (|qGI| > 0.6, FDR < 1%) GIs is indicated in each precision and recall heatmap. **(h)** Replicate agreement of genome-wide qGI scores (n = 17,804) of selected pairs of the 7 query genes that were replicated multiple times. The Pearson’s correlation coefficient (PCC) r, and the WBC score (see Methods) quantify qGI reproducibility. The WBC score is a z-score-based metric that indicates how much the Pearson correlation coefficient (PCC) between a pair of qGI profiles differs from the average PCC observed across different genetic backgrounds (see Methods). **(i)** Mean WBC score for seven genetic backgrounds at different normalization stages of the qGI pipeline. The mean per genetic background was taken across all within-background pairs for *FASN* (n = 4), *PDCD5* (n = 4), *FANCG* (n = 4), *PTAR1* (n = 4), *NGLY1* (n = 4), *PELO* (n = 5) and *VPS52* (n = 5). A WBC score is a z-score-based metric, and the mean WBC for each genetic background passes the threshold of three only after complete normalization. **(j)** Mean replicate correlation coefficient for seven genetic backgrounds at different normalization stages of the qGI pipeline.

### Reproducibility analysis of qGI-derived interactions

In addition to evaluating the impact of the qGI normalization components, we also performed a comprehensive characterization of the reproducibility of the qGIs measured by our pipeline. To do so, we generated a comprehensive dataset comprising seven diverse query backgrounds and, in total, 30 biologically replicated, genome-wide query mutant screens. We applied a novel MCMC-based method, called JEDER^62,69,70^, on this data to estimate the precision and recall of GIs identified. From the four to five independent screens performed for each of the queries, JEDER generates a GI consensus profile, which can then be used to determine precision and recall at different combinations of thresholds based on interaction effect size, the qGI, and the associated FDR. We selected two separate qGI/FDR threshold pairs to provide “standard” or “stringent” GI calls, which reflect different balances in the tradeoff between precision and recall (Fig. 6f, g). Standard GIs were defined at a JEDER-estimated precision of 61% and a recall of 37%, which was achieved at an absolute qGI score > 0.3 and an FDR < 10%. Stringent GIs were defined at a precision of 91% and a recall of 12%, which was achieved at |qGI| > 0.6 and FDR < 1%. Those two threshold pairs robustly identify GIs from genome-wide CRISPR-Cas9 screens in the context of various experimental artifacts as well as empirically defined reproducibility metrics.

In addition to reproducibility of thresholded interactions, we evaluated the quantitative reproducibility of interaction profiles. Depending on the number of significant interactions, which varied broadly across the different query backgrounds we screened, between-replicate Pearson’s correlation coefficients of genome-wide qGI scores ranged from approximately 0.2 for the *PTAR1* query screens to 0.71 for the *VPS52* screens (Fig. 6h, j; Fig. S7g). Using the Within-versus-Between-context replicate Correlation (WBC) score, a metric displaying how well a pair of replicates is distinguished from screens performed in another genetic background that can be interpreted as a z-score^71^, we found that all replicates passed the threshold of WBC > 3. Between-replicate correlation coefficients consistently decreased at every correction step, but importantly, the distinction of replicates from different genetic backgrounds, as measured by the WBC score, increased substantially as steps of the qGI pipeline were applied (Fig. 6i, j; Fig. S7f). This suggests that experimental artifacts present within data produced by genome-wide CRISPR screens can contribute to non-specific background correlation between biologically unrelated screens, and that our qGI normalization pipeline can effectively remove those artifacts.

## Discussion

We report the qGI score, a computational pipeline for accurate identification and quantification of GIs from genome-wide CRISPR-Cas9 screens. Our framework incorporates a series of data correction steps to control for experimental artifacts, including genomic linkage effects and biologically driven, GI-independent covariation between screens. The identification of these artifacts was unexpected given the use of query mutants derived from a single cell line and underscores the value of systematic evaluation of controlled experimental systems. Additionally, we applied an MCMC-based approach to empirically characterize the precision and recall of qGI-derived interactions of different effect sizes and FDR thresholds, applied across a large panel of replicated screens. Collectively, we provide a detailed and general scoring pipeline for and analyses of GIs derived from genome-wide CRISPR screens. We anticipate that both the insights about technical artifacts and the correction strategies will be broadly applicable to other differential, or context-specific, CRISPR screening studies including chemical-genetic screens.

Context-specific CRISPR screens, where the context reflects a specific cell line, cell type or tissue, can be strongly affected by false positive (FP) differential fitness effects resulting from genomic linkage, via at least two different well-described mechanisms^50,51,53,54,63^. Most prominently, unequal DNA copy number (CN) between cell lines can exacerbate fitness effects for genes located within CN-altered regions^54^. CRISPR screens derived from query mutants in a single cell line are designed to avoid this by introducing a precise mutation in an otherwise identical genome. However, our analysis revealed chromosomal regions exhibiting systematic biases in GI scores, resulting in numerous FP, and to a lesser extent, false negative GIs, as well as linkage-driven biases in GI profile analyses.

We found evidence for multiple mechanisms underlying these genomic linkage artifacts. First, local CN alterations where a duplicated region in a query clone is compared to a unique region in the control produce spurious positive GIs for many genes in the duplicated region. This phenomenon parallels the CN effect previously described for context-specific essential genes, suggesting that CN effects should be considered when comparing query mutants to their wild-type control cell line. In another scenario, strong negative GIs involving a library gene located near a chromosome end, such as *TUBB4B* in the *TUBB* knockout clone, produced numerous, spurious negative GIs. As described by Lazar and colleagues in the Cancer Dependency Map (DepMap)^63^, sporadic loss of the chromosome arm may cause collateral loss of the strong interacting gene. Despite evidence supporting the presence of CN or chromosome arm loss effects in some cases, those mechanisms explain only a minority of the genomic linkage artifacts we observed. Importantly, we found that large-scale GI biases affecting entire chromosomes, especially chrX, occurred in nearly one-third of query screens, suggesting that epigenetic regulation of sex chromosomes may require specific attention.

Context-specific functional screening datasets, such as our HAP1 GI network and the DepMap network, are typically generated over several years, during which experiments are conducted in different batches and under varying conditions. Batch effects and random noise are well-recognized challenges and can be addressed through established normalization strategies^9,13,53,54^, which are implemented in our qGI pipeline. However, beyond technical variability, we identified pervasive, biologically coherent covariation across both control and replicated query screens, with effect sizes comparable to the strongest GIs in a typical genome-wide query screen. Traditional genomics quality control approaches typically check for biological coherent signal among top effects using functional enrichment tests or related approaches. Such techniques would be blind to FP GIs resulting from the between-screen covariation patterns we describe here, and may even identify spurious signal as evidence of strong screen quality. To correct for these effects, we developed an SVD-based method that identifies and removes shared patterns of variation present in HAP1 control screens from each query screen, in a screen-specific manner. This correction will have a substantial impact on the analyses of context-specific screens, because it deprioritizes numerous genes central to highly investigated research areas, such as cancer biology and aging. We found these genes to vary strongly and frequently across control screens in coherent patterns. Importantly, the effectiveness of this correction depended on our large set of replicated control screens, which enabled us to estimate the systematic variation introduced by our CRISPR screening platform. We expect that this approach could be generally applied to interpret context-specific functional screening data and improve the utility of these experiments as powerful discovery tools.

In the qGI scoring pipeline, correction modules sequentially reduced between-replicate screen correlation while progressively increasing query-specific signal reproducibility, as measured by the within-vs-between-replicate correlation (WBC) score. These improvements were reflected by increased functional information derived from the resulting GI profiles following each correction step. Together, the correction modules integrated into the qGI pipeline markedly improve the accuracy and specificity of GI quantification from genome-wide CRISPR screens and provide a means for accurate, comprehensive GI analysis in human cells.

## Methods

### Pooled genome-wide CRISPR-Cas9 dropout screening in HAP1 query mutant cell lines

Human HAP1 wild type cells were obtained from Horizon Genomics (clone C631, sex: male with lost Y chromosome, RRID: CVCL_Y019). HAP1 gene knockout cell lines were either obtained from Horizon or generated in this study. See Billmann *et al.* for cell line annotation files and experimental details^62^. All gene knockout cell lines were confirmed to carry the expected out-of-frame insertions or deletions.

The experimental steps were previously described^32,72^. In brief, CRISPR library virus production was performed in HEK293T cells using the lentiviral lentiCRISPRv2 vector containing the TKOv3 gRNA library (Addgene #90294)^72^. Virus-containing media was harvested 48 hours after transfection and aliquoted and frozen at -80°C.

For determination of viral titers for pooled screens in HAP1 cells, multiplicity of infection (MOI) of the titrated virus was determined 72 hours post-infection by comparing percent survival of puromycin-selected cells to infected but non-selected control cells.

Pooled genome-wide screens were performed as previously described^32,34^. A total of 90 million HAP1 cells were transduced with lentivirus containing the TKOv3 gRNA library at a MOI of ∼0.3, such that each gRNA was represented in ∼200-300 cells. Transduced cells were allowed to recover for 24 h, selected using puromycin (1ug/mL) for 48 hours, and then a pellet was collected to represent the T0 starting population. The remaining cells were split into triplicates, and cultured for up to 20 population doublings while maintaining a library coverage of at least 200-fold throughout. Cells were passaged every 3-4 days and cell pellets, one from each triplicate, were collected to represent the midpoint and endpoint populations. Screens were performed in either DMEM with 10mM glucose and 1mM glutamine (‘minimal medium’), or IMDM with 25mM glucose and 1mM pyruvate (‘rich medium’) (see Billmann *et al.* for data file^62^). All media was supplemented with 10% FBS and 1% penicillin–streptomycin, and cultures were grown at 37°C with 5% CO2.

The qGI score was developed and characterized using an initial set of 363 genome-wide screens (324 query mutant + 39 WT controls) all of which passed basic quality control such as separation of core essential and non-essential genes. For final genetic interaction network analysis, as described^62^, a subset of the query mutant screens was removed based on manual inspection and more detailed QC filters as detailed in Billmann *el al.*^62^, which resulted in a total set of 298 query mutant screens used in that study.

### Illumina sequencing and mapping and quantification of gRNA reads

From the collected cell pellets, the library preparation for sequencing was performed as described previously ^32^. In brief, sequencing libraries were prepared from 50 µg of the extracted genomic DNA in two PCR steps, the first to enrich guide RNA regions from the genome, and the second to amplify guide-RNA and attach Illumina TruSeq adapters with i5 and i7 indices as described previously using staggered primers aligning in both orientations to the guide-RNA region (see Aregger *et al.*^32^ for details). Barcoded libraries were gel purified and final concentrations were estimated by qRT-PCR. Sequencing libraries were sequenced on an Illumina HiSeq2500 using single read sequencing and completed with standard primers for dual indexing with HiSeq SBS Kit v4 reagents. The first 21 cycles of sequencing were dark cycles, or base additions without imaging. The actual 36-bases read begins after the dark cycles and contains two index reads, reading the i7 first, followed by i5 sequences. The T0 and T18 time point samples were sequenced at 400- and 200-fold library coverage, respectively (Fig. S1b, f).

FASTQ files from single read sequencing runs were first trimmed by locating constant sequence anchors and extracting the 20 bp gRNA sequence preceding the anchor sequence. Pre-processed reads were aligned to a FASTA file containing the TKOv3 library sequences using Bowtie (v0.12.8) allowing up to 2 mismatches and 1 exact alignment (specific parameters: -v2 - m1 -p4 --sam-nohead).

### Estimation of fitness effects in HAP1 cells

To quantify genetic interactions (GIs), the fitness of HAP1 cells was recorded upon single and combinatorial perturbation of genes. In the following sections, CRISPR-Cas9-based gene perturbations are referred to as “mutants”. Single mutant fitness (SMF) effects were estimated for each individual control and query screen and for each gRNA first. This was done by computing the log_2_ fold-changes (LFC) between read-depth normalized gRNA abundance of gRNAs in the TKOv3 library^72^ in the starting population (T0) and the endpoint (10-12 population doublings) (T18) to quantify each gRNA’s dropout from the pool at the end of the experiment:

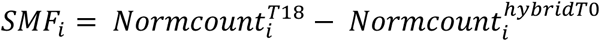

where hybrid T0 refers to a matched T0 for each control and query screen that is additionally stabilized (defined in detail in the next section). The read-depth of gRNAs in all samples, T0 and T18, was normalized using the following equation:

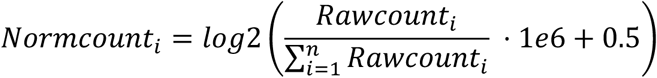

Where 𝑖 represents a gRNA in the library, and 𝑛 is the total number of gRNA in the library.

After read-depth normalization, gRNAs with a readcount before normalization below 40 or above 10,000 in a T0 sample were excluded from further analyses (Fig. S1c). The upper threshold was used to control extreme but very rare outliers. Of the 71,090 gRNAs in the library 2,092 (1,050 on the (-), 1042 on the (+) strand) gRNAs were flagged in at least one experiment. Of those, 985 gRNAs, including 367 gRNAs that were not in the library, were flagged in at least 20% of the T0 samples (Fig. S1e), and were removed from any sample.

Population doublings were determined by cell counting throughout the course of the experiment and depended on media conditions and query mutations. Throughout the process of completing the screens for this study, several plasmids containing the TKOv3 library and lentiviral packaged plasmids were required, which introduced dependencies between batches of screens. Moreover, we observed that gRNA abundance after library infection and puromycin selection differed from the abundance of the library, and that this difference was highly reproducible. Those T0 gRNA populations showed a tendency towards virus batch-driven patterns. This suggested that effects on the measurement of the starting population throughout the duration it took to record all CRISPR-Cas9 screens for the GIN would be manifested in the SMF effects if not accounted for during the LFC estimation. In such cases, query-specific effects could be driven by the abundance changes before and after library infection. The qGI score uses matched T0 measurement to assure that differences between screens during library infection and Puromycin selection would not result in false positive (FP) GIs. In addition, we implemented a matched T0 stabilization by also using the median across all 𝑚 T0 measurements of the same virus batch (common T0). Those two estimates were combined in a weighted fashion to minimize correlation between GI scores and the difference between matched and common T0 readcounts:

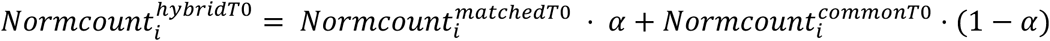

If the number 𝑚 of genome-wide screens in a given batch is odd,

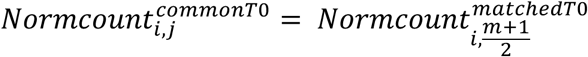

If the number 𝑚 of genome-wide screens in a given batch is even,

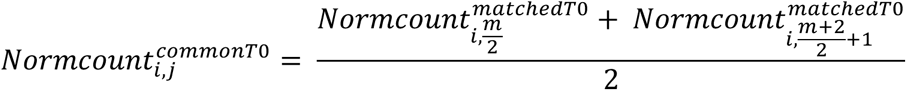

Where 𝑖 represents a TKOv3 library gRNA in a given genome-wide screen 𝑗, and 𝑚 represents the number of genome-wide screens conducted in the same batch as 𝑗.

𝑁𝑜𝑟𝑚𝑐𝑜𝑢𝑛𝑡*^matchedTo^* and 𝑁𝑜𝑟𝑚𝑐𝑜𝑢𝑛𝑡*^commoTo^* were balanced to compute the

𝑁𝑜𝑟𝑚𝑐𝑜𝑢𝑛𝑡*^hybridTo^* by finding the optimal 𝛼, where 0 ≤ 𝛼 ≤ 1:

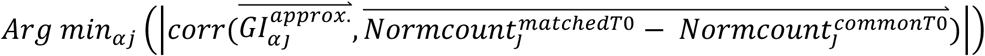

𝛼 was sampled 31 times in increments of 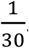. To speed up the scoring process, gRNA-level GI scores were approximated by sampling gRNAs as well as estimating the GI Null model more efficiently.

Finally, for each control WT and query mutant genome-wide screen, the SMF effects were adjusted, by setting the median of non-essential genes, as defined in Hart *et al.* ^20^, to 0.

### Estimation of residual fitness effects

To score GIs, SMF effects estimated using the genome-wide TKOv3 gRNA library were compared between an isogenic HAP1 cell line harbouring a stable query mutation and the parental WT HAP1 cell line. To obtain robust estimates of SMF effects in the two media conditions used for screening, 21 and 18 genome-wide screens were performed in WT HAP1 cell lines cultured in minimal and rich media, respectively. In the following, we will refer to those 21 and 18 control screens as WT “control” screens.

The TKOv3 library contains 71,090 guide (g)RNAs that target about 18k human protein-coding genes, most of them with four sequence-independent gRNAs. gRNA-level residual scores were derived for a given genetic background by estimating a non-interacting model between LFC values in this background and each control screen. For each WT-KO screen pair, the population of LFC values were M-A-transformed ^73^, which contrasts the per-gRNA LFC difference (M) with per-gRNA arithmetic mean (A). Since SMF values are represented by log2-foldchanges (LFC), we computed M and A as follows:

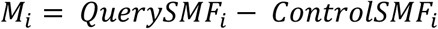

and

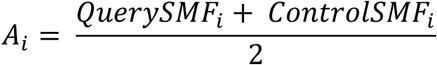

where *i* represents each gRNA, ranging from 1 to 71,090. To obtain the null model for each query screen, a second-degree Loess regression for each KO-WT screen pair was least squares-fit along 𝐴 using the R stats function loess. A was additionally locally stabilized by binning the data along 𝐴 with equal numbers of data points in every bin to result in the vectors 𝑀*_balanced_* and 𝐴*_balanced_*.

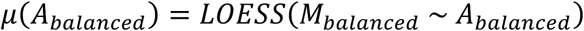

Where 𝜇(𝐴) is the smooth regression function estimated from bin-balanced 𝑀*_balanced_* and 𝐴*_balanced_* and represents the genetic interaction null model. 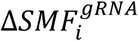 represents the per-gRNA residual SMF effects, which are the gRNA-level interaction scores:

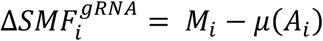

For each gRNA, this resulted in 21 (minimal media) or 18 (rich media) residual SMF effects, which represent the contrasts of a given KO with the 21 or 18 WT HAP1 control screens, respectively.

Overall, the qGI score compares for each screening condition all query screens to all control WT screens. In addition, for each condition, all control screens are contrasted amongst each other to identify pseudo-(FP) GI effects. For the first GIN cell culture condition in minimal media, we contrasted 21 control screens to each other and 147 query screens. For rich media conditions, we contrasted 18 control screens to each other and 177 query screens. The control screens were also contrasted against earlier sampling time points of the control screens to cover a wide spectrum of screen sampling time points when modeling and correcting for GI-independent screen-to-screen variation and covariation (see below). Together, this resulted in 7524 pairwise comparisons between genome-wide screens including 1251 comparisons representing pseudo-GI effects. We noted that the range of residual effects differed substantially, and that this variation was largely driven by SMF dynamic ranges and not associated with query phenotypes. Therefore, the qGI score scales residual SMF effects per screen pair by adjusting the standard deviation of the central 80% of gRNA residual values. Under the assumption that specific query screens can have extreme interaction effects, the top 10% of most extreme positive and negative gRNA residual SMF phenotypes are not considered for standard deviation estimation. Scaling of residual SMF effects globally improved the ability of gene-level qGI scores to recapitulate genes’ co-complex or co-pathway membership.

### Robust estimation of library gene single mutant fitness (SMF) effects

Every query screen is contrasted to multiple control WT HAP1 screens, which provides a stable estimate of the differential SMF effects caused by the query mutation. At the same time, control screens that share the same putative (unknown) batch effect as a given query would have fewer differential SMF effects that those with a different set of batch effects. Since putative batch effects of screens are unknown, we applied an unsupervised weighting of all query-control screen pairs. The weights were assigned based on the assumption that genetic interactions are sparse^74^. We reasoned that experimental artifacts such as batch effects would introduce additional signal into the population of residual values and assigned a higher weight to WT HAP1 screen contrasts with lower sum of squared residuals mean across the 95% least extreme ∼70,000 gRNAs in the library. Excluding the top 2.5% most negative and most positive SMF residuals, respectively, prevented penalizing WT-query contrasts with a small number of very strong residuals, which could, in contrast to consistently large residuals, represent true GIs. Based on those weights, all 21 query-control pairs in minimal and all 18 query-control pairs in rich media were summarized, resulting in a single ∼70,000 gRNA-long vector of stabilized residual SMF effects for each query. We refer to the resulting value for each gRNA as the “guide-level” GI score.

### Correction of genome position-linked biases of residual SMF effects

On-target CRISPR-Cas9-introduced DNA cuts are accompanied by target gene-unspecific DNA repair-associated toxicity, which has been shown to associate with copy number (CN) and genomic position and can create false essential gene calls in CRISPR screens if uncorrected^54^. Supervised and unsupervised techniques to correct for falsely identified essential genes are routinely applied ^51,53,54^. Those methods are applied on SMF data, and unequal FP SMF effects in control and query screens can be expected to cause FP GIs. In our context, given that we are performing isogenic screens in which one query gene is perturbed in a common HAP1 parental background, we reasoned that any CN or other chromosomal artifacts should largely be shared between the WT and mutant HAP1 cells. Thus, any chromosomal artifacts present uniquely in the query would become evident in the residual SMF effects (e.g., contrasting the query mutant screen with the WT control screens). We designed our chromosomal artifact correction focusing on these residual SMF effects and developed an algorithm that to find such genome position-linked biases of residual SMF effects.

Biases are identified for residual SMF effects at the gRNA-level. As described above, those residual SMF effects were robustly estimated by taking the weighted consensus of a given query screen compared to all WT control HAP1 screens in the same cell culture condition. This assured that any detected bias can be attributed to the query screen (instead of a control screen).

First, the qGI pipeline tests for differences between mean residual SMF effects 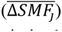 of adjacent groups of gRNA values while moving along genome coordinates, where *j* is in *1* to 71,090 − 𝑛. The mean residual SMF effects are defined as:

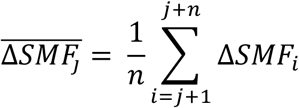

With this mean residual SMF effect, differences between groups are tested as:

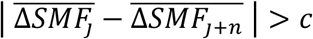

The size of this group is defined per chromosome and is higher for chromosomes for a higher gRNA per bp ratio. The group size thus covers comparable distances of the genome while maintaining a number of measurements (gRNA values) that allows a stable estimate of the residual SMF effect. The group size depended on the number of gRNA on each chromosome 𝑘_0%(_ (see R code implementation for details).

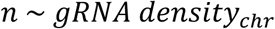

If a difference between mean residual SMF effects (dLFC) of adjacent groups of gRNA values is detected, its genome coordinate (X) is saved. Biases in residual SMF effects between X_i_ and X_i+1_ or, for the first and last X on a chromosome, between X_i_ and the start or end of the chromosome are tested. If the mean residual SMF effects in such a genomic region deviates from 0 (positive and negative deviation is possible) more than a defined threshold (flexible parameter, set at > 0.15 for this study), this region is saved as a putatively biased region.

Finally, to avoid affecting between-gRNA variance for genes at the boundaries of detected regions, gRNAs were assigned to a region if all gRNAs targeting the same gene were identified. Those regions are sub-chromosomal. In addition, potential residual SMF biases affecting each entire chromosome are identified for chromosomes without detectable local biases. For this whole-chromosome test, we measured if the mean gRNA-level residual SMF effect for the entire chromosome deviated from 0 (positively or negatively) more than a defined threshold (flexible parameter, set at >0.09 for this study). For any regions with identified genome position-linked biases, the residual SMF effects were corrected by subtracting the mean effect from all gRNA-level measurements in the affected region.

### Unsupervised correction of GI-independent interaction patterns using SVD

The qGI score was designed to robustly identify GIs, which are differential SMF effects between a query and WT control screen that can be attributed to the query mutation. A simple comparison between control and query CRISPR screens is obscured by a compendium of factors contributing to structured screen-to-screen variability, many of which are poorly understood. In our data, screen-to-screen variability affects both the control as well as the query screens (Fig. 4a-i; Fig. S6a-d). The qGI score implements a series of correction steps that estimate artifacts from the large collection of control and query mutant data and robustly correct for those artifacts. Unwanted screen-to-screen variance as well as GI-independent covariance patterns are removed via a two-step singular value decomposition (SVD)-based qGI correction module.

In the first step, the qGI scoring pipeline estimates and corrects screen-to-screen variation patterns in screen-to-screen differential SMF effects from WT control screens. Specifically, like a query screen, each WT control screen is contrasted to all remaining control screens in the same media condition. Again, as for the query screens, all resulting contrasts for a given control screen with all other control screens are summarized. This generates a matrix of control screens (n = 21 for minimal media; n = 18 for rich media) X gRNAs (∼70,000) of residual SMF effects, only that residual SMF effects are not the result of a query mutation. This captures all GI-independent (i.e. no query gene) screen-to-screen variation. In addition, the effective number of doublings past library infection when a screen is stopped can affect SMF phenotypes ^60,67^, which we refer to as temporal screen-to-screen variation, and this can occur when query cell lines grow substantially slower than HAP1 WT cells and the cell population goes through substantially fewer cell doublings. Therefore, additional intermediate time points between day 6 and day 17 were collected and like all individual endpoint control screens, are contrasted to the remaining endpoint control screens. This causes strong residual SMF effects (Fig. 4b) and captures temporal screen-to-screen variation. This extended the residual SMF effects matrix to 40 (21 of which are endpoint) and 25 (18 of which are endpoint) WT control screens times gRNAs (∼70,000) for minimal and rich media, respectively. Utilizing these control screen data matrices, the query screen residual SMF effect matrices are corrected in multiple steps.

First, the GI-independent residual SMF effect variance in query data was estimated for each library gRNA. This was done by computing the per gRNA-variance across query screens as well as control screens. Here, intermediate time point data was not considered, because those were taken from a screen that was represented by an endpoint and samples taken from the same screen are highly correlated, which impacts variance estimates. For the entire gRNA population, the query variance was fitted via a first-degree loess model with a span of 0.5. This per-gRNA estimate was then scaled to have a mean of 1 and query screen as well as all control screen residual SMF effects were divided by this variance estimate.

Second, GI-independent covariance patterns were removed. This was done by decomposing the 40 (min) or 25 (rich) control screens 𝑐 × ∼70,000 gRNA 𝑙 variance-corrected residual fitness effect matrices via singular value decomposition (SVD) using the R function svd(). NAs were imputed using row mean values. The left singular vectors 𝐿*_l×c_* were then multiplied to create a matrix 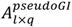.

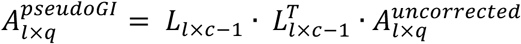

The unit matrix was projected onto the variance-corrected query residual SMF effect matrix, and the result was then subtracted from the variance-corrected query residual SMF effect matrix.

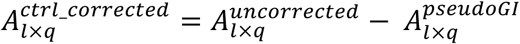

This removal of GI-independent covariation patterns was done for each media condition separately.

In the second GI-independent pattern normalization step, the qGI scoring pipeline removes dominant query screen covariation patterns. Previous studies involving large genetic screens have demonstrated the utility of removing dominant covariation patterns to purify underlying sparse, biological signal^6,13,53,68^. The qGI pipeline decomposes the corrected residual SMF effect matrix between all non-redundant query screens and the ∼70,000 gRNAs via SVD. To avoid capturing latent variables that represent GI screens with a larger number of biological replicates, query genes with larger numbers of biological replicate screens were represented by one of those replicates. Just as for the control WT screens, NAs were imputed using row mean values. The left singular vectors 𝐿_=×DJ_ were then multiplied to create a matrix 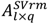.

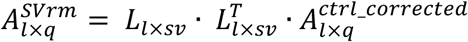

This was done by setting the first, the first two, first three, etc. singular vectors to 0. The different matrices were projected onto the query residual fitness effect matrix 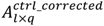 and the resulting representation 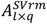 was subtracted from the query residual SMF effect matrix.

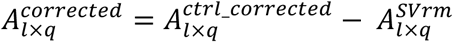

This gRNA-level GI score (corrected residual SMF effect) matrix was summarized at the gene level and the capacity of GI profiles to reconstruct CORUM 3.0 co-complex^75^, co-pathway (MSigDB Bioconductor package) and co-GO biological process memberships was assessed using the FLEX pipeline ^67^. Based on the precision-recall performance, the first four singular vectors were removed from the data presented in this study. The decision of how many singular vectors should be removed is a dataset-dependent decision that should be made with all available information about screens and an objective benchmarking pipeline (e.g. FLEX) ^67^.

### Empirical TKOv3 library gRNA quality score (QS)

Before summarizing gRNA-level into gene-level qGI scores and computing the associated FDR, gRNAs were tested for their gene-specific phenotype, and poorly performing gRNAs were excluded based on an empirical TKOv3 gRNA quality score (gRNA QS). This gRNA QS is based on the observation that interaction profiles are highly informative of gene function and the assumption that independent gRNA sequences targeting the same gene have similar interaction profiles. We further assumed that the majority of independent gRNA sequences for each gene are very unlikely to have off-target signal that is correlated, and that any observed strong GI profile correlation between gRNAs targeting the same gene is on target. Therefore, an outlier gRNA that does not correlate with the other gRNAs against the same gene, when others do correlate, is assumed to perform poorly and should not be used for computing gene-level scores.

To find and exclude those poorly performing gRNAs, for each gene targeted by the TKOv3 library, the pairwise Pearson’s correlation coefficient (PCC) was computed between all gRNA targeting that gene along the corrected gRNA-level GI scores across all 324 query screens. For all gRNAs targeting the same gene, the mean of PCCs was computed (excluding self-correlation), resulting in the gRNA QS. Based on this gRNA QS, each gene was tested for outlier gRNAs. First, gRNAs were only allowed to be flagged if other gRNAs targeting the same gene showed evidence of coherent interaction signal because we reasoned that reducing the number of gRNAs for genes with subtle interaction signal is likely to magnify spurious interaction signal driven by a single gRNA. This was judged based on the assumption that true and detectable interaction signal of at least two gRNAs targeting the same gene pass a minimum gRNA QS. The threshold was chosen such that 40% of the genes (7,129 of 17,804 genes) were considered to have sufficient interaction signal based on the mostly positive within-gene gRNA interaction score correlation (Fig. S2c). Selecting a higher threshold further increases this positive shift for within-gene correlations and would increase the confidence in gRNA QS-based calls but would also limit the sensitivity to flag low-quality gRNAs. The large majority of the selected genes show evidence of expression in HAP1 (Fig. S2d, e), but a substantial portion of these genes show no significant SMF effect (∼40.7% are not detected at FDR < 20%) (Fig. S2f).

Second, for each of those 7,129 genes with reliable interaction signal, the worst gRNA per-gene based on its QS was flagged if it had a large negative differential gRNA QS to the per-gene median gRNA QS by a second threshold shortlisting the worst gRNAs for 777 (4.4% of 17804) genes. Amongst this set of implicated gRNAs, a subset was kept if their gRNA QS surpassed a third threshold showing reasonable agreement with the other gRNA against the same gene, resulting in a reduced set of 517 genes (2.9% of 17804) flagged for low gRNA QS. For those 517 genes, the worst gRNA was removed before computing the SMF, the qGI score and the qGI-associated FDR (Fig. S2g).

### Gene-level quantitative genetic interaction (qGI) scores

After the corrections described above were applied, we computed a quantitative, gene-level score summarizing the GI effect per query gene-library gene combination. The qGI score consists of an effect size measure (a corrected differential log2-fold change) as well as an associated per-screen false discovery rate (FDR), representing the statistical confidence in a non-zero interaction for that gene pair, accounting for the multiple tests per screen. Gene-level GI scores are computed by mean-summarizing corrected gRNA values, with poor empirical TKOv3 gRNA library quality scores removed. Therefore, the resulting qGI score benefits from all previously described correction steps. The qGI-associated FDR is computed based on residual SMF effects that are corrected for coarse screen-specific effects (i.e., genomic linkage, residual-on-readcount abundance dependency, per-screen residual SMF effect scaling, WT-query contrast weighting) but without between-screen normalization steps (i.e., SVD-based corrections for screen-to-screen variance and covariance). Also, the multiple control WT contrasts for each query screen were not summarized before estimating FDR. For each query-library gene pair, a *p*-value was computed using limma’s moderated *t*-test, which considered independent gRNAs targeting the same library gene as biological replicates and values derived from different control vs. query screen comparisons as technical replicates^76^. Here, limma’s duplicateCorrelation() function is used to estimate the inter-duplicate correlation for each query screen (Fig. S2a, b). For each query screen, *p*-values were corrected for multiple hypothesis testing using the Benjamini-Hochberg method, resulting in the qGI-associated q-values, which we refer to hereafter as the FDR.

### Evaluation of GI reproducibility

To quantify qGI score reproducibility, the <u>W</u>ithin-vs.-<u>B</u>etween context replicate <u>C</u>orrelation (WBC) score (described in ^71^) was computed on all query screens with two or more biological replicates. To summarize briefly, first, the pairwise Pearson’s correlation coefficient was computed between all biological replicate query screen pairs. As described in Billmann et al. ^62,71^, biological replicate correlations were then scaled to the expected background correlation distribution: the mean and standard deviation of its correlation with screens performed in another context (e.g., query mutation). This converted the correlation coefficients into a metric with an unambiguous statistical interpretation that can be interpreted as a *z*-score. Here, each query creates its own background correlation distribution. If all biological replicates for a given query were performed in the same media condition, only all other query screens in this media condition were used as background. If two biological replicates were performed in different media conditions, the background was only generated across media conditions (e.g., the minimal media replicate obtained its background correlation distribution from all rich media screens and vice versa). To judge all biological replicates for a given query gene separately, as well as all screens sharing the same query gene as a whole, WBC scores were computed for all possible pairs as well as collectively for the entire set of replicates for a given query. Note that the selection of query screens used for creating each query screen’s own background correlation distribution determines the exact numerical WBC score, and scores for an identical set of replicates for a given query will not be identical in a growing data set where the background correlation distribution is estimated from more queries.

### Functional evaluation of GIs

To functionally evaluate GIs, we tested how well GI profiles across all 324 query screens in our data set recapitulated known biology as provided by various data bases. Specifically, we assumed that a positive Pearson’s correlation coefficient (PCC) between the qGI profiles along the 324 query screens of a gene pair would be indicative of functional similarity. We utilized the FLEX package to test how well our data reconstructed CORUM 3.0 co-complex^75^, co-pathway (MSigDB Bioconductor package) and co-functional (GO biological process) annotations ^67^.

### Analysis of genomic linkage of SMF effects in the DepMap

We tested if genome position-linked GI scores in the HAP1 isogenic cell lines set were indicative of a general trend beyond HAP1 cells. For chromosomal and sub-chromosomal regions with recurrent residual SMF effect biases in HAP1 cells, we also quantified the density of strong co-essentiality in the Cancer Dependency Map (DepMap)^77^. Chronos gene SMF scores (version 23Q2) were downloaded from https://depmap.org/portal/. The data contained 1,095 genome-wide CRISPR-Cas9 gene knockout screens with scores corrected for proximity bias^50^. To test the relative local co-essentiality density, co-essentiality was computed as the PCC along the Chronos score profiles across 1,095 cell lines. The ratio of the density of edges with a z-score larger than four between genes in a given region over the density among all genes was computed.

---

**Figure S1:**
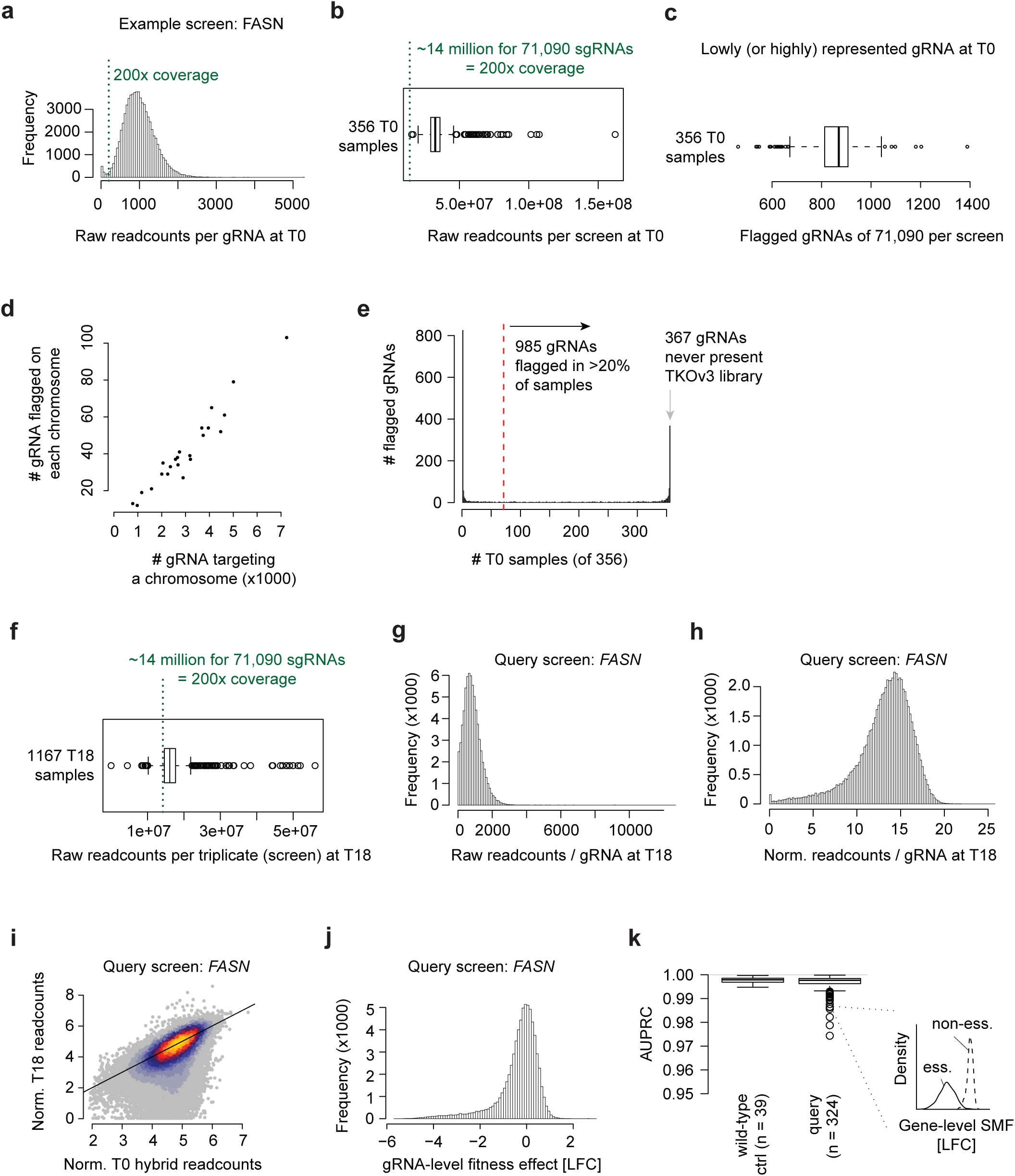
Initial quality control of genome-wide CRISPR-Cas9 screens. **(a)** Raw readcounts before sequencing depth normalization per gRNA for all 71,090 gRNAs in the TKOv3 library at T0. Shown is a FASN query screen as a representative example. **(b)** Total number of raw readcounts per screen at T0 for each query and control screen. Overall, 356 unique T0 samples from the 363 screens existed in the initial dataset, since several control screens that were performed in different media conditions shared a common T0. **(c)** Number of gRNAs with readcounts before read-depth normalization below 40 for each screen. **(d)** Mean number of gRNAs flagged on each chromosome compared to the number of gRNAs targeting genes on those chromosomes. **(e)** Number of screens a gRNA is flagged in. Only the 2092 gRNAs that are flagged in at least one T0 sample (screen) are shown. **(f)** Total number of raw readcounts per screen at T18 (screen endpoint) for each query and control screen. Overall, 1167 unique end and intermediate time point samples (technical triplicates) were sequenced from the 363 unique screens. **(g, h)** Raw (g) and normalized log2 (h) readcounts per gRNA for all 71,090 gRNAs in the TKOv3 library at T18 (endpoint). Shown is a FASN query screen as a representative example. **(i)** Normalized log2 readcount T0 hybrid versus T18 (endpoint) comparison of all 71,090 gRNAs in the TKOv3 library. Shown is a FASN query screen as a representative example. **(j)** T18 over T0 hybrid foldchange (lfc) of the FASN screen. **(k)** Area under the precision-recall curve (AUPRC) distinguishing the Hart *et al.* core essential genes from non-essential genes as a basic per-screen quality control statistic.

**Figure S2:**
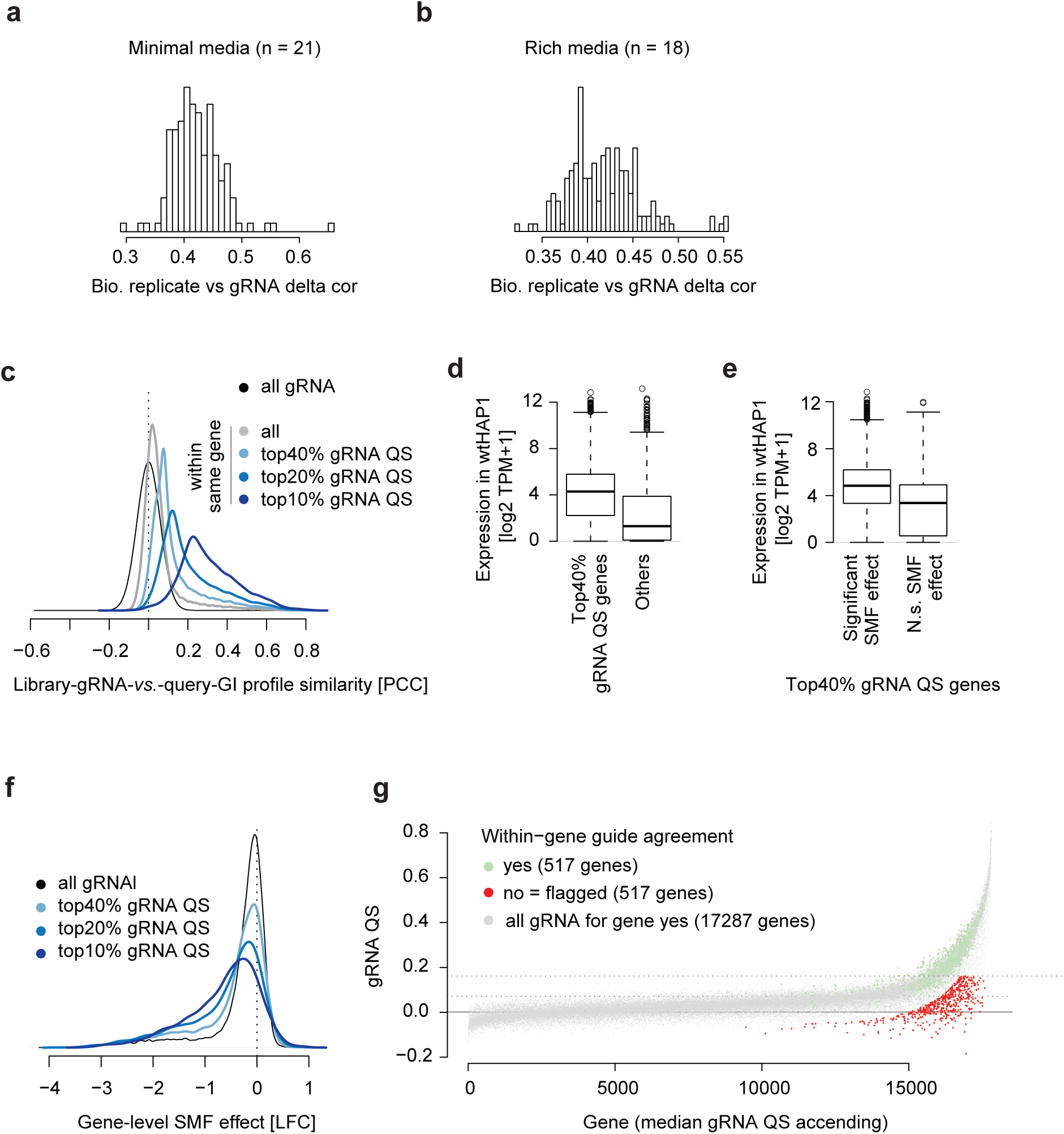
Agreement and quality control of gRNAs targeting the same gene. (a, **b)** Relative replicate correlation of gRNA-level GI scores to determine the setup of technical and biological replicates in limma’s moderated t-test. This is computed independently for the minimal and rich media screening conditions. **(c)** GI profile similarity of all pairwise comparisons of gRNAs targeting the same gene. **(d)** Expression of the genes with the top 40% gRNA QS in HAP1 WT cells. **(e)** Expression of the genes with the top 40% gRNA QS separated by whether they showed a single mutant fitness effect at a lenient FDR of 20% in HAP1 WT cells. **(f)** single mutant fitness (LFC) for different groups of top gRNA QS genes. **(g)** The gRNA QS for all ∼70,000 gRNAs grouped by targeted gene. Flagged gRNAs are highlighted in red.

**Figure S3:**
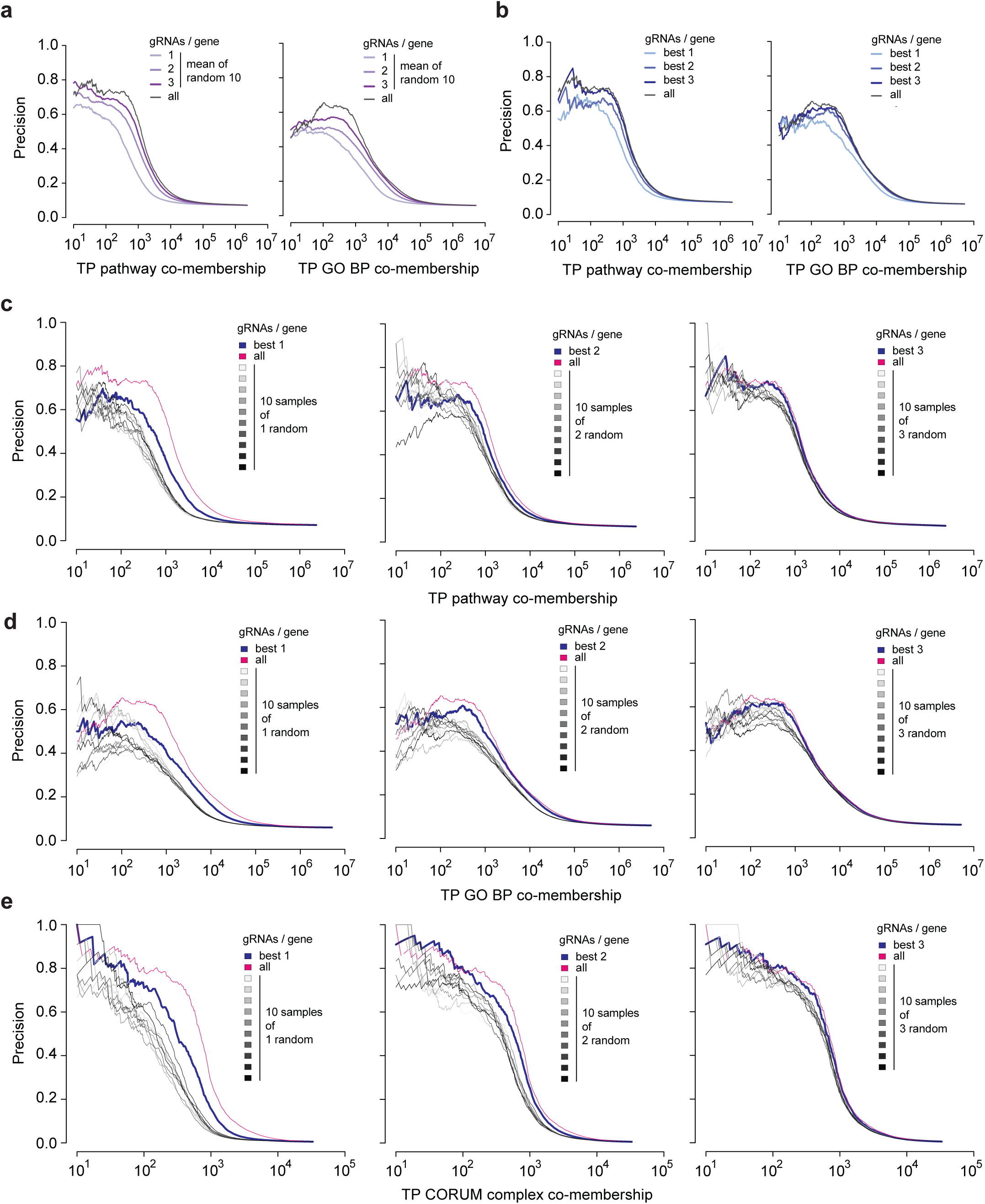
Effect of gRNAs-per gene on performance of qGI-based profiles to identify complex, pathway or GO-BP co-membership. qGI scores are based on 1, 2, 3 or all up to 4 gRNAs per gene. qGI profile Pearson’s correlation coefficients across 324 query screens are used as a similarity metric. **(a)** Mean precision of 10 iterations of 1, 2 or 3 randomly selected gRNAs targeting a specific gene compared to all up to 4 gRNAs per gene. Shown is the performance of for identifying pathway or GO BO co-membership. Corresponds to Fig. 2g. **(b)** Top 1, 2 or 3 gRNAs per gene with the best gRNA QS score compared to all up to 4 gRNAs per gene. Shown is the performance of for identifying pathway or GO BO co-membership. Corresponds to Fig. 2h. **(c-e)** Comparison of the best 1, 2 or 3 gRNAs targeting a specific with 10 iterations of 1, 2 or 3 randomly selected gRNAs.

**Figure S4:**
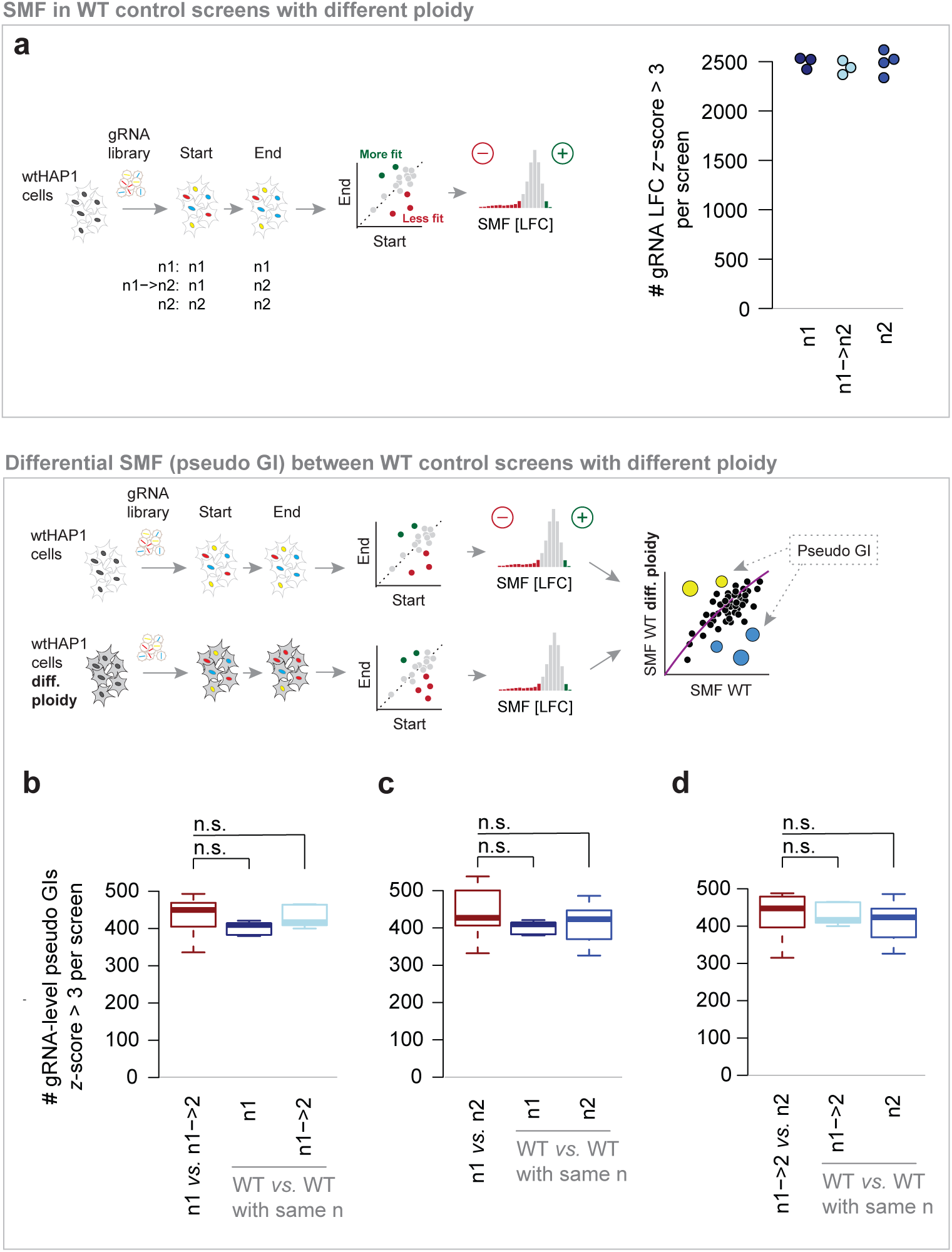
Number of differential SMF effects between WT HAP1 screens with different ploidy status. **(a)** Number of strong fitness effects (gRNA-level LFC z-score < -3) in screens with 1n at both start and end of the experiment, screens that developed from 1n to 2n, and those that were 2n from the start. Each dot represents a single genome-wide screen. **(b-d)** Number of strong differential effects (gRNA-level uncorrected interaction z-score < -3) when screens with different ploidy status were contrasted (red). No statistical difference was observed when comparing to the number of strong differential effects is between screens with the same ploidy (Wilcoxon’s rank sum test).

**Figure S5:**
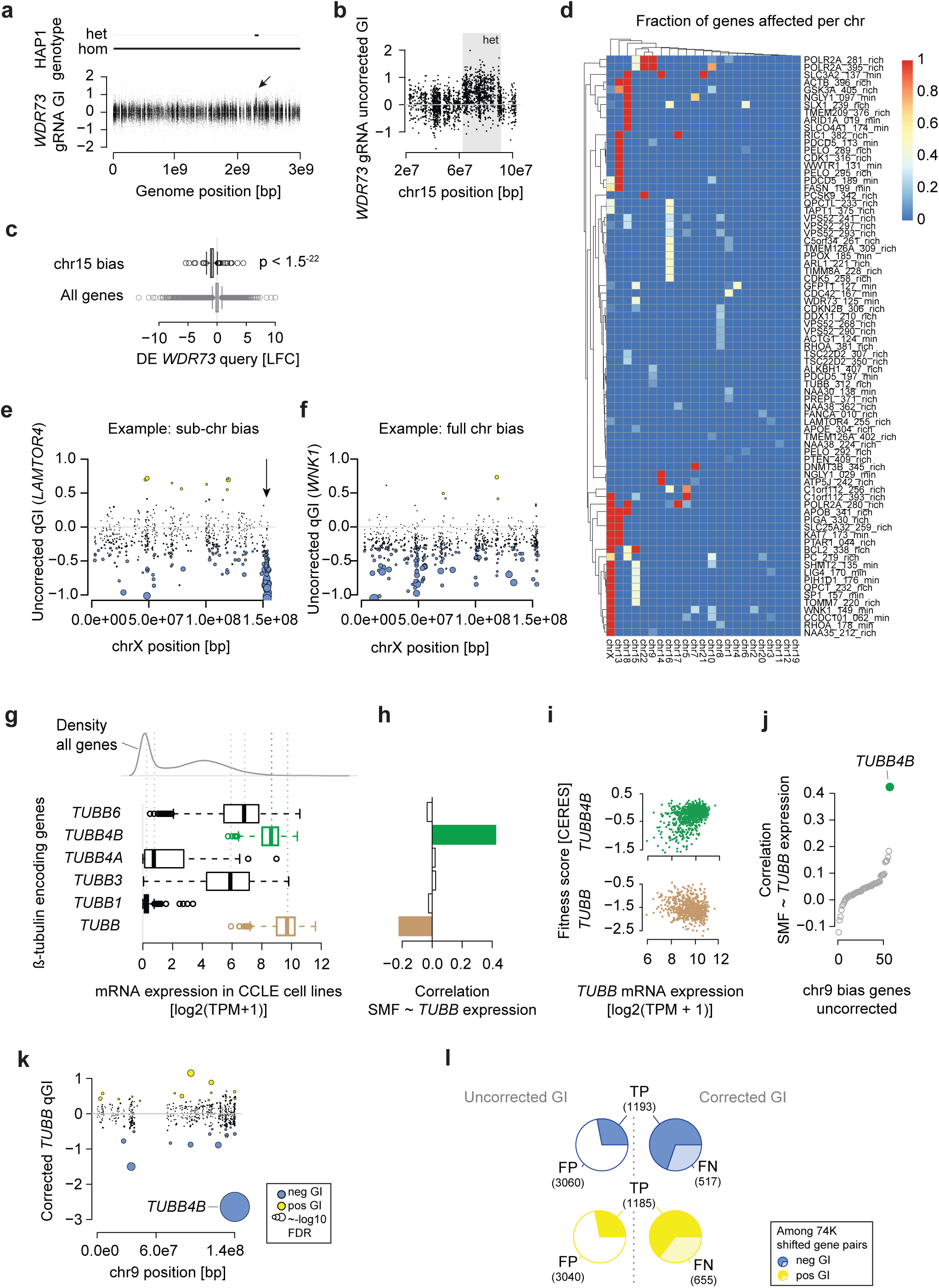
Exploration of genome position-linked false positive genetic interactions. **(a)** gRNA-level, uncorrected qGI scores showing interactions with a *WDR73* mutation in the HAP1 cell line are plotted against the genome position of their target sequence. The heterozygous 30 Mbp region on chr15 in parental HAP1 cells is shown and the arrow points at uncorrected gRNA-level GI scores in this region. **(b)** Uncorrected *WDR73* gRNA-level qGI scores on chr15 (zoomed in view of region highlighted in (a)). **(c)** Differential expression distributions in the *WDR73* KO clone of all genes in the genome compared to the genes on in the heterozygous chr15 region. **(d)** Fraction of genes per chromosome with detected biases. Shown are query screens with at least one affected chromosome other than chrX. **(e, f)** Uncorrected *LAMTOR4* and *WNK1* qGI scores on chrX. Dot size corresponds to -log10 FDR of the qGI score, and colors indicate significant GIs (|qGI| > 0.3, FDR < 0.1). **(g)** Expression of φ3-tubulin encoding genes in the Cancer Cell Line Encyclopedia. **(h, i)** Correlation between *TUBB* expression and fitness effects upon perturbation of beta tubulin encoding genes in the Cancer Dependency Map (DepMap). **(j)** Correlation between *TUBB* expression and DepMap fitness effects of the genes in the affected chr9 q34 region. **(k)** Corrected *TUBB* qGI scores on chr9. Dot size corresponds to -log10 FDR of the qGI score, and colors indicate significant GIs (|qGI| > 0.3, FDR < 0.1). **(l)** Number of positive (yellow) and negative (blue) GIs (|qGI| > 0.3, FDR < 0.1) across all genome regions with detected bias in the 324 screens that were gained (FN), removed (FP) and retained (TP) after correction.

**Figure S6:**
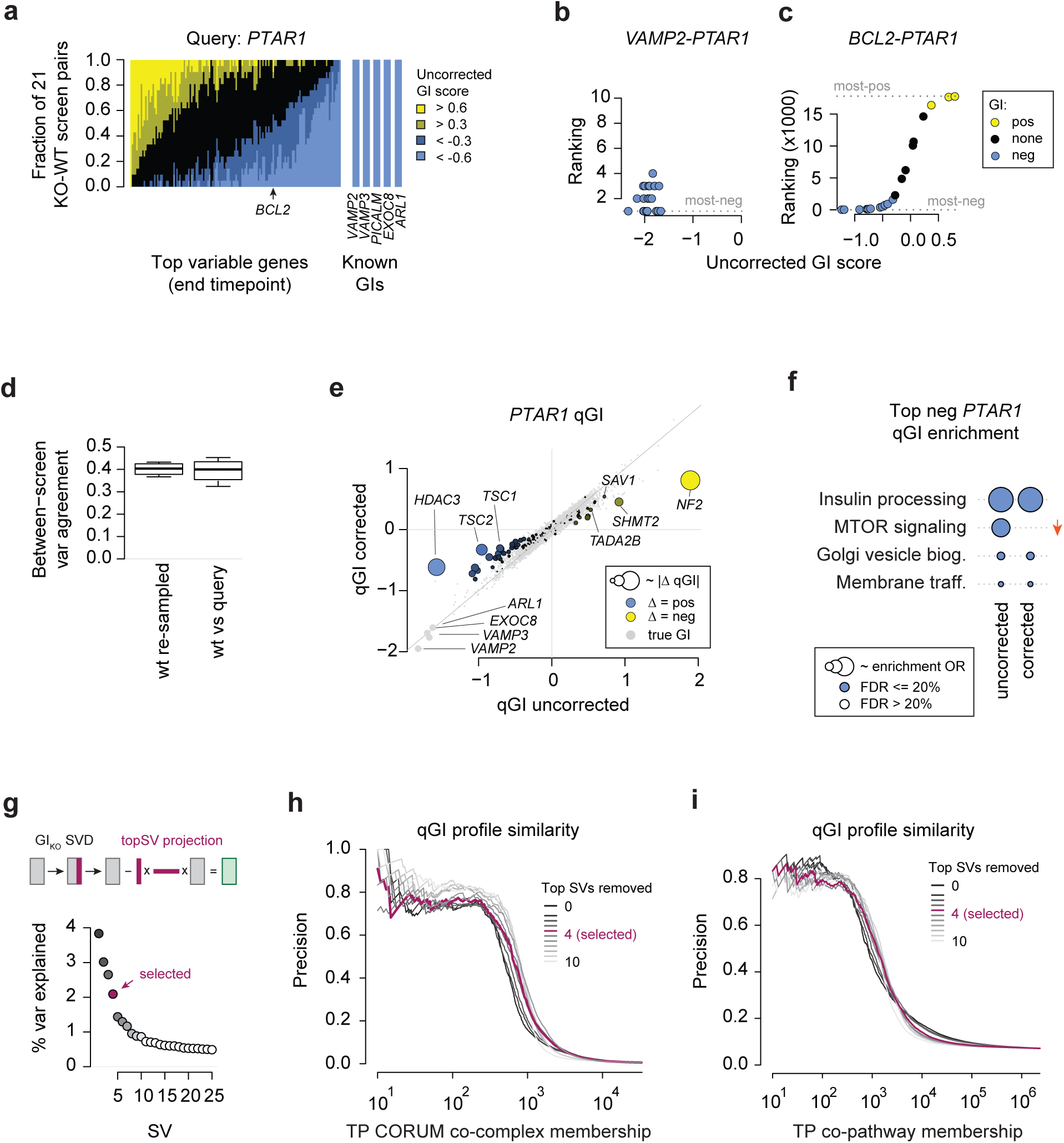
Unsupervised correction of genetic interaction-independent patterns in genetic interaction screens. **(a)** Fraction of positive and negative uncorrected qGI scores of highly variable genes and 5 high confidence negative GIs with *FASN* in all 21 KO-WT contrasts. The ∼140 highly variable genes were selected because they exhibited the highest fitness effect-adjusted variance between control screens or exhibited high covariance in the control screens. **(b, c)** Ranking and qGI scores (uncorrected) between *PTAR1* and *VAMP2* (known) or the highly variable gene *BCL2* in the 21 WT contrasts with three independent *FASN* KO screens. **(d)** Between-biological replicate variance agreement of WT control and query screens. The standard deviation across the residual fitness scores of biological replicated screens was computed for each library gene. This scaled representation of the between-screen variance was compared between *FASN*, *FANCG*, *NGLY1* and *PDCD5* query screen sets and the WT control. **(e)** *PTAR1* qGI scores before and after correction for genetic interaction-independent patterns. Shown are 17,804 genes, and the highly variable genes are highlighted. Dot size corresponds to the gene-specific change of the qGI score. A correction towards a more positive value (less negative qGI score) is shown in blue, and correction toward more negative (less positive qGI score) in yellow. **(f)** REACTOME pathway enrichment of the top 100 negative and positive qGI scores in the PTAR1 query screen before and after correction. All pathways enriched at an FDR of 20% before or after are shown. **(g)** Variance explained by the first 25 singular vectors (SV). Singular vector decomposition (SVD) was performed on the gRNA-level qGI matrix and considering every unique query genetic background (see Methods). **(h, i)** Precision-recall performance of qGI profiles with increased number of SVs removed to reconstruct known co-complex (h) or co-pathway (i) memberships. Co-complex memberships were defined by CORUM 3.0, co-pathway memberships were defined by MSigDB. Top SVs were cumulatively removed from the data. Based on these performance evaluations, the version removing SV1-4 was selected.

**Figure S7:**
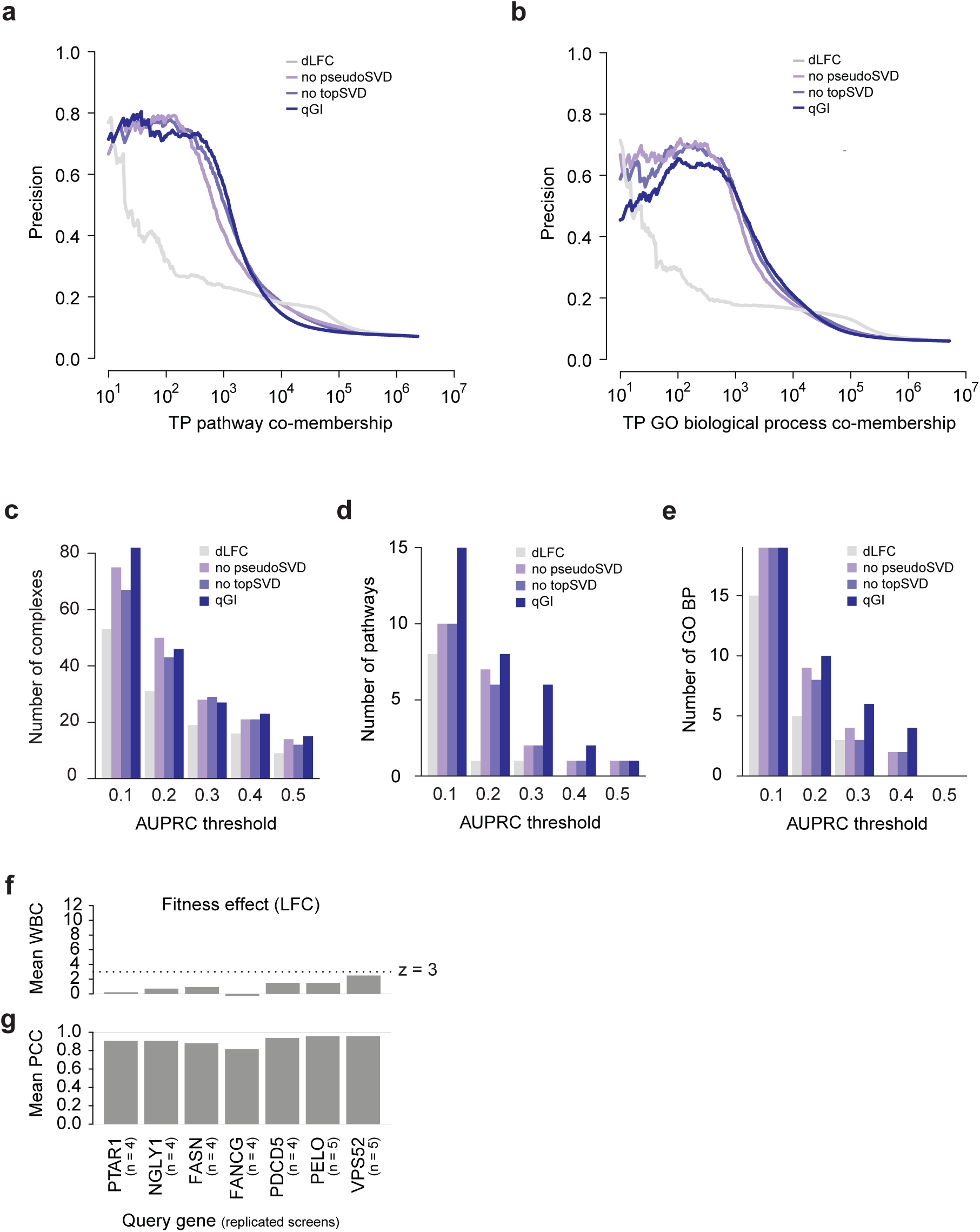
Performance of qGI profiles at different normalization stages of the qGI pipeline to reconstruct known complex, pathway and GO-BP co-memberships. Shown is the simple differential (d)LFC, a fitted raw qGI version without removal of GI-independent patterns and the overall top four SVs, a version without removal of the overall top four SVs, and the completely corrected qGI score. qGI profile Pearson’s correlation coefficients across 324 query screens are used as a similarity metric. **(a, b)** Precision-TP curve for pathway (a) and GO BP (b) as functional standard. This figure corresponds to Fig. 6a. **(c-e)** Number of CORUM complexes (c), pathways (d) and GO BPs (e) different module-level AUPRC thresholds. Redundancy of modules (complexes, pathways, GO BPs) is controlled at a Jaccard index of 0.8. **(f)** Mean WBC score for seven genetic backgrounds of the fitness scores (LFC). The mean per genetic background was taken across all within-background pairs for *FASN* (n = 4), *PDCD5* (n = 4), *FANCG* (n = 4), *PTAR1* (n = 4), *NGLY1* (n = 4), *PELO* (n = 5) and *VPS52* (n = 5). A WBC score is a z-score-based metric. **(g)** Mean replicate correlation coefficient for seven genetic backgrounds of the SMF scores (LFC).

